# Predicting the retinotopic organization of human visual cortex from anatomy using geometric deep learning

**DOI:** 10.1101/2020.02.11.934471

**Authors:** Fernanda L. Ribeiro, Steffen Bollmann, Alexander M. Puckett

## Abstract

Whether it be in a single neuron or a more complex biological system like the human brain, form and function are often directly related. The functional organization of human visual cortex, for instance, is tightly coupled with the underlying anatomy with cortical shape having been shown to be a useful predictor of the retinotopic organization in early visual cortex. Although the current state-of-the-art in predicting retinotopic maps is able to account for gross individual differences, such models are unable to account for any idiosyncratic differences in the structure-function relationship from anatomical information alone due to their initial assumption of a template. Here we developed a geometric deep learning model capable of exploiting the actual structure of the cortex to learn the complex relationship between brain function and anatomy in human visual cortex such that more realistic and idiosyncratic maps could be predicted. We show that our neural network was not only able to predict the functional organization throughout the visual cortical hierarchy, but that it was also able to predict nuanced variations across individuals. Although we demonstrate its utility for modeling the relationship between structure and function in human visual cortex, our approach is flexible and well-suited for a range of other applications involving data structured in non-Euclidean spaces.

## 1 Introduction

The modern-day study of visual perception has led to an unprecedented understanding of the neurobiological mechanisms underlying sensory information processing in the brain. Although first being of interest to those pursuing basic science, this understanding has since translated to more practical applications, notably by inspiring the development of convolutional neural networks (CNNs) (Fukushima, 1980) which have revolutionized the field of artificial intelligence (Lecun et al., 2015). In the human visual system, light reaches the retina where it is converted into electrical impulses that are transferred to the lateral geniculate nucleus and then through a hierarchy of cortical areas (Figure 1) (Felleman and Van Essen, 1991; Hubel and Wiesel, 1962; Mishkin et al., 1983). At the bottom level of the hierarchy (primary visual cortex), neurons with small receptive fields (i.e., the coverage area in the visual field to which a neuron is responsive) capture simple visual features such as spatial orientation (Hubel and Wiesel, 1962). As those signals travel through higher order visual areas (comprised of neurons with increasing receptive field sizes) those simple visual features are combined, enabling complex features to be mapped from the visual input. These more complex features range from contours and textures to complete faces near the top of the hierarchy (Kanwisher et al., 1997). This type of hierarchical processing is imitated by CNNs by using filters with learnable parameters which are convolved across data representations with the aim of capturing common features in the input (Seeliger et al., 2018). At early layers of CNNs, those filters capture simple features, while in deeper layers more complex features are detected due to the complex combination of the inputs of previous layers. Here, we report that the story has – in a sense – come full circle. By leveraging recent advances in the application of CNNs to non-Euclidean spaces (Bronstein et al., 2017), we show that deep learning can be used to predict the functional organization of the very brain regions that inspired them (i.e., the areas of the visual cortical hierarchy).

**Figure 1.**
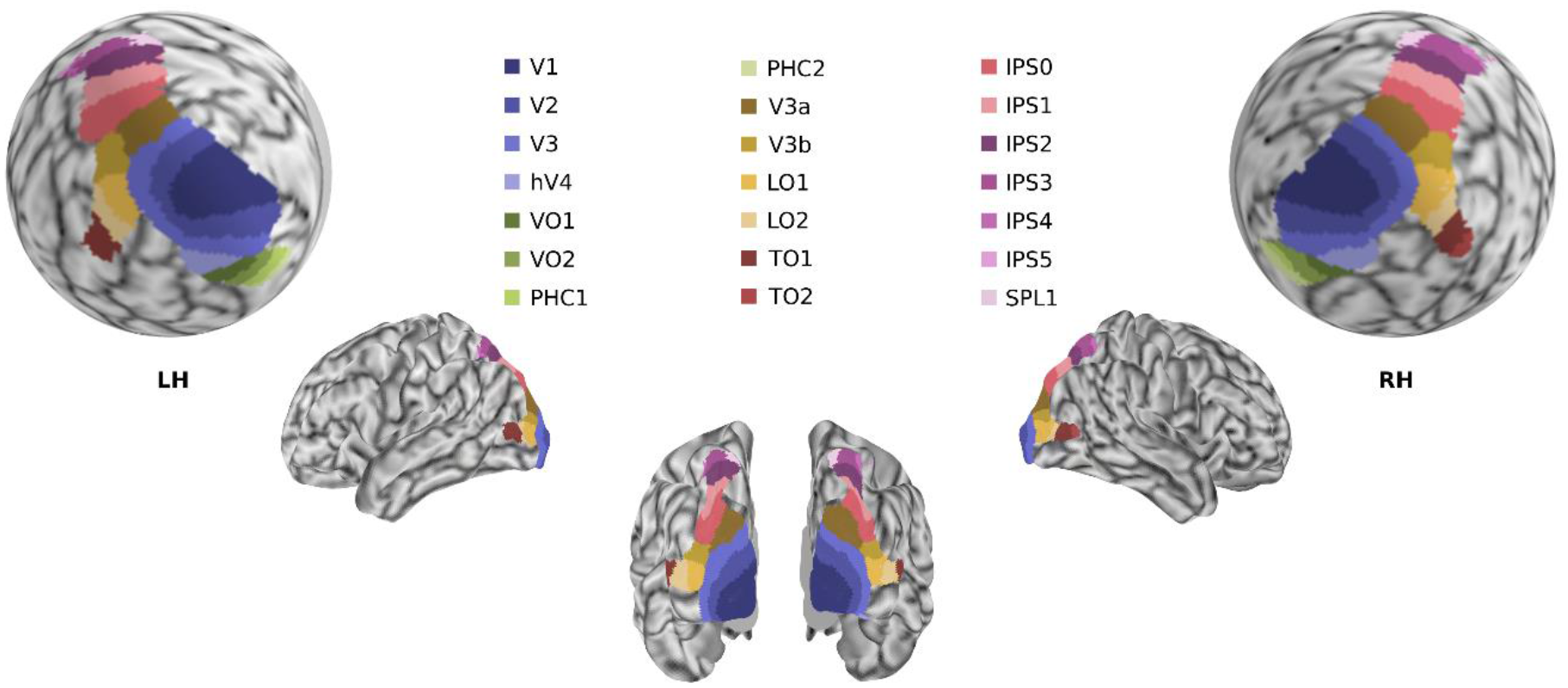
Hierarchical organization of the visual cortex. The 21 atlas-based (Wang et al., 2015) visual areas in both left (LH) and right (RH) hemispheres displayed on spherical surface and fiducial surface (lateral and posterior views) models (Van Essen et al., 2013). Different colors represent different visual areas. Central legend indicates the name of each visual area.

The visual hierarchy is comprised of a number of functionally specialized cortical visual areas (Zeki et al., 1991), nearly all of which are organized retinotopically (Grill-Spector and Malach, 2004; Van Essen, 2004). That is, the spatial organization of the retina (and hence the visual field) is maintained in each of these cortical visual areas - albeit in a distorted fashion due to properties such as cortical magnification (i.e., there are more neurons dedicated to processing foveal vs. peripheral information) (Cohen, 2011; Cowey and Rolls, 1974). In other words, adjacent neurons in each of these retinotopically organized visual areas map adjacent objects in the visual field. Although distorted, this retinotopic mapping is known to be similar across individuals and the mapping functions between the visual field and the brain, at least in early visual cortex, can be described by relatively simple mathematical models (Balasubramanian et al., 2002; Schira et al., 2007). However, considerable inter-individual variation does exist (Van Essen and Glasser, 2018), and this variation has been shown to be directly related to variability in gross anatomical properties, such as variability in brain size, cortical folding patterns, and the degree of myelination (Abdollahi et al., 2014; Benson et al., 2014, 2012; Benson and Winawer, 2018; Hinds et al., 2009; Rajimehr and Tootell, 2009; Wu et al., 2012). Note, for instance, that the size of primary visual cortex (V1) and hence its corresponding retinotopic maps vary by a factor of 2-3 between individuals (Dougherty et al., 2003).

Given the variability in retinotopically organized cortex across individuals, it has previously been held that obtaining an accurate estimate of a particular individual’s retinotopic maps requires a dedicated empirical experiment. These experiments, whether done using a conventional phase-encoded design (DeYoe et al., 1996; Engel et al., 1997; Sereno et al., 1995) or a more modern population receptive field (pRF) approach (Carvalho et al., 2020; Dumoulin and Wandell, 2008; Zeidman et al., 2018), are capable of producing robust and detailed estimates of these maps. Both of these approaches entail stimulating different visual field locations while an individual undergoes an fMRI scan – and then subsequently examining the elicited cortical activation with respect to the known spatiotemporal properties of the visual stimulus. These empirically derived maps can be used specifically to examine the functional organization of the visual cortex or for more general purposes such as defining visual area boundaries, which is instrumental for any visual experiment aiming to isolate or compare cortical area specific signals. Despite being a well-established and reliable approach, collecting the retinotopic mapping data needed to construct these empirical maps still requires significant resources (e.g., scan time is usually between 15-60 minutes) and can be impractical or even impossible in certain circumstances (e.g., some clinical populations). As such, considerable effort has been dedicated to developing an alternative method for estimating these maps at the individual level.

As mentioned, the functional organization of the visual hierarchy is tightly associated with its underlying structure. Given that structural data is easier and less time consuming (requiring only 5-15 minutes) to collect than functional data, the notion of predicting the retinotopic maps directly from the underlying structure has been attractive – and cortical shape has, in fact, been shown to be a useful predictor of retinotopic organization in early visual cortex (Benson et al., 2014, 2012; Hinds et al., 2009; Schira et al., 2012; Wang et al., 2015). The current state-of-the-art in predicting retinotopic maps uses mathematical templates that are warped to an individual’s cortical anatomy and is able to account for gross individual differences in the functional organization of early visual cortex (Benson et al., 2014, 2012). While becoming widely used instead of empirical retinotopic mapping and of clear utility (Albers et al., 2018; Alvarez et al., 2019; Isherwood et al., 2017; Storrs et al., 2020), such models are unable to account for any idiosyncratic differences in the structure-function relationship from anatomical information alone – due to their initial assumption of a template. Here, it was our aim to develop a predictive model capable of capturing the intricate structure-function relationship of the visual cortex without enforcing a spatially consistent mapping (i.e., a template), such that more realistic and idiosyncratic maps could be predicted. To this end, we set out to train a CNN using the most comprehensive and detailed retinotopy dataset openly available – that from the Human Connectome Project (HCP) (Benson et al., 2018). Note that this dataset includes ultra-high field (7T) fMRI retinotopic mapping data of a considerable number of participants (181 participants) along with their anatomical data, serving as an ideal dataset for this endeavor.

Convolutional neural networks are conventionally applied to data represented in two- or three-dimensional Euclidean space (Figure 2a). A great deal of data, however, is not best represented in such a simple and regular domain (Bronstein et al., 2017). In the field of neuroscience, and for retinotopic mapping in particular, data are often represented on surface models of the brain as many cortical properties only make sense considering their specific location with respect to the various sulci and gyri (Van Essen et al., 1998). Indeed, the HCP 7T retinotopy dataset being used here was made available with the functional and structural data already mapped onto surface-based representations of the cortex (Benson et al., 2018). These surface models are essentially sheet-like meshes that are intricately folded in three-dimensional space, maintaining the geometrical structure of the brain (Borne et al., 2020; Dale et al., 1999; Van Essen and Glasser, 2018). Importantly, two points on the surface model might be close to one another in Euclidean space but far apart in terms of cortical distance following the surface (e.g., on opposing banks in a sulcus). To account for this and other non-Euclidean properties, techniques have recently been developed that generalize deep neural network models to non-Euclidean spaces such as surfaces (Figure 2b) and graphs – with these techniques collectively being referred to as geometric deep learning (Bronstein et al., 2017).

**Figure 2.**
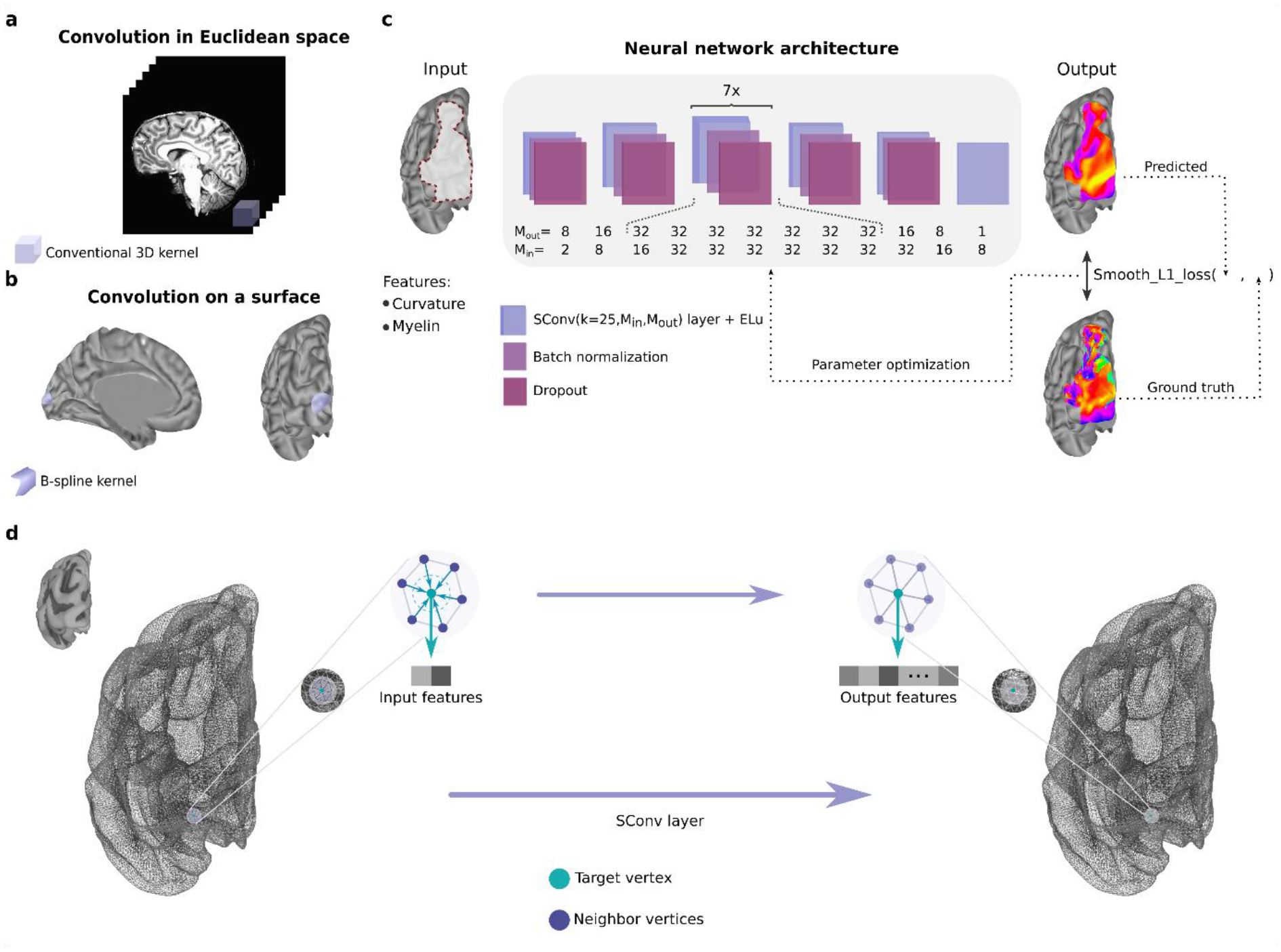
Predicting brain function with geometric deep learning. **a,** Convolution in 3D Euclidean space involves convolving a conventional 3D kernel across multiple slices of 2D images. **b,** Convolution on a surface, a non-Euclidean space, involves convolving a B-spline kernel directly on the surface, in which the kernel is able to aggregate information in local neighborhoods. **c**, Geometric convolutional neural network architecture. Input data included curvature and myelin values (vertex features) as well as the connectivity among the ROI surface vertices (i.e., the surface topology) and their spatial disposition (i.e., their 3D coordinates) in a cortical surface (implicit feature). The output data is the vertex-wise polar angle/eccentricity values. Network parameters were optimized to reduce the difference between prediction and ground truth using a smooth L1 loss function. **d**, SConv layers aggregate vertex features in local neighborhoods weighted by learnable parameters of a continuous kernel function (Fey et al., 2018). This process generates new attributes for each target vertex at a time. Thus, while the cortical surface space was the same for all individuals, allowing for vertex correspondence across individuals, vertex features varied.

Here, we leveraged these recent advances in machine learning to achieve our aim of developing a framework capable of learning the complex relationship between the functional organization of visual cortex and the underlying anatomy using the surface-based HCP 7T retinotopy dataset. That is, using geometric deep learning we developed a neural network capable of predicting the retinotopic organization of the human visual hierarchy (Figure 2c). Our model was not only able to predict the functional organization of human visual cortex from anatomical properties alone, but remarkably it was also able to predict nuanced variations across individuals. Although we show its utility for retinotopic mapping, geometric deep learning is flexible and well-suited for an abundance of other applications in the field of neuroscience (Gopinath et al., 2018; Seong et al., 2018; Zhao et al., 2019).

## 2 Results

### 2.1 Retinotopic mapping with deep learning

Retinotopic mapping typically involves empirical data collection performed by showing visual stimuli, which are temporally and spatially controlled, to a participant lying in an MRI scanner (DeYoe et al., 1996; Dumoulin and Wandell, 2008). Analysis of the fMRI signals elicited by such stimuli allows the visual field location encoded by each responsive voxel to be estimated. The visual field locations are usually defined in polar coordinates resulting in two retinotopic maps: one representing the polar angle (or clock position) and the other representing the eccentricity (or distance away from fixation). An example of these empirically derived retinotopic maps for a single individual is shown in Figure 3 (top row). From these, it can be seen that the polar angle and eccentricity maps are organized roughly orthogonal to one another – most easily discerned in early visual cortex (V1, V2, and V3). Since the human visual system is comprised of multiple retinotopically organized cortical areas, these maps are also used to demarcate the boundaries of the various areas. The polar angle maps are particularly useful for this as the boundaries between areas can be identified by reversals in the progression of the polar angle values. In Figure 3, for example, the superior boundary between V1 and V2 is found by locating the band at which the polar angle reverses from progressing from red to yellow, back to red. Likewise the inferior boundary between V1 and V2 is found where the polar angle representation reverses from progressing from red to purple, back to red.

**Figure 3.**
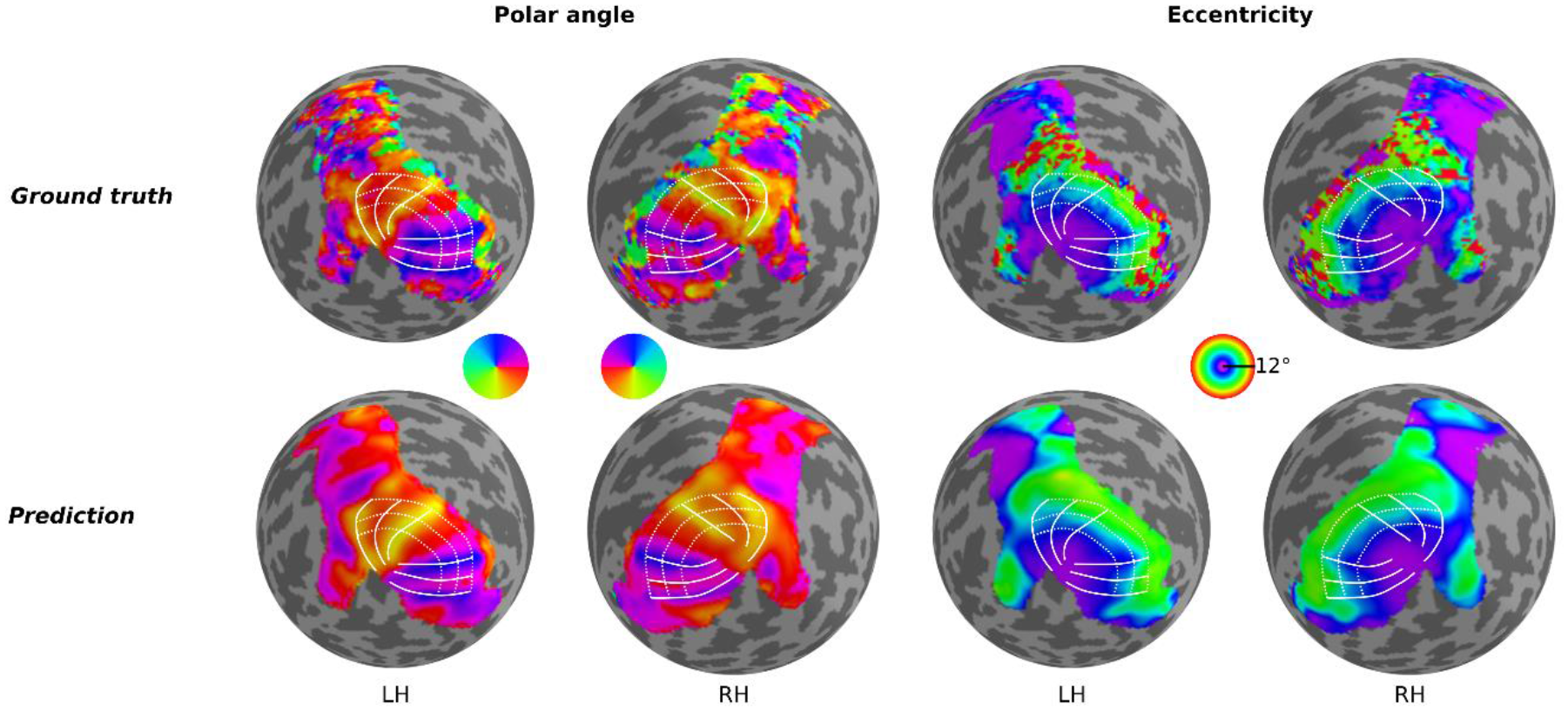
Retinotopic mapping with geometric deep learning. Upper row shows empirical (ground truth) polar angle and eccentricity maps for both left (LH) and right (RH) hemispheres of a single individual from the test dataset (*Participant 1*). Lower row shows the predicted polar angle and eccentricity maps for each hemisphere of the same participant. Polar angles vary from 0⁰ to 360⁰, while eccentricity values varied from 0⁰ to 12⁰ as indicated by the color wheels. The white grid illustrates isoeccentricity (dashed) and isopolar angle (solid) lines associated with early visual cortex (V1, V2, and V3). Note that V2 and V3 are split around V1 and are comprised of dorsal and ventral components.

To map retinotopically organized areas without the need to collect functional data, we built a geometric convolutional neural network by stacking spline-based convolution layers (Fey et al., 2018) to compute convolutions on a manifold (i.e., the cortical surface model) (Figure 2c). The data serving as input to the network included curvature and myelin (determined by the ratio of T1w/T2w images) values resampled to the HCP 32k fs_LR standard space, as well as the connectivity among vertices forming the surface (i.e., the surface topology) and their spatial disposition (i.e., their 3D coordinates) in the HCP 32k fs_LR standard surface space (Figure 2d). The output of the model was either the polar angle or eccentricity value for each vertex of the cortical surface model. Developing the neural network involved three main steps: (1) *training* the neural network, (2) hyperparameter *tuning*, and (3) *testing* the model. Prior to the training step, the 181 participants from the HCP dataset were randomly separated into three datasets: training (161 participants), development (10 participants) and test (10 participants). These datasets were used in each of the above three steps, respectively. During *training*, the network learned the correspondence between the retinotopic maps and the anatomical features by exposing the network to each example in the training dataset. Hence, the empirically-derived retinotopic maps served as the ground truth and the parameters of the neural network were optimized to minimize the difference between the predicted and empirical retinotopic maps. Note that the objective function, smooth L1 loss, was weighted by the explained variance from the population receptive field (pRF) modeling procedure used to generate the empirical retinotopic maps (Benson et al., 2018; see section 4.6 in the Methods). Model hyperparameters, such as the number of layers, were *tuned* (i.e., optimized) by inspecting model performance using the development dataset. Finally, once the final model was selected, the network was *tested* by assessing the predicted maps for each participant in the test dataset. This procedure was followed for each hemisphere and each type of retinotopic mapping data separately, resulting in four predictive models.

Figure 3 illustrates the predicted retinotopic maps with their corresponding ground truth (i.e., empirically measured retinotopic maps) for a single participant in the test dataset. Qualitatively, the predicted maps bear a remarkable resemblance to the ground truth, appearing to accurately predict the main features of both polar angle and eccentricity maps. To aid comparison between the empirical and predicted maps a grid of isopolar angle and isoeccentricity lines has been overlaid upon the maps in early visual cortex. The grid was drawn based on the ground truth data and then positioned identically on the predicted maps. Note how well the visual area boundaries (i.e., the isopolar angle contours) match the predicted maps, running directly through the polar angle reversals. A similar quality of fit can be seen for the isoeccentricity lines.

Although the grid is focused on early visual areas (due to their organization being most well understood and hence most reliable to outline), it is important to note that the neural networks are able to predict the retinotopic organization far beyond early visual cortex, throughout the visual hierarchy (Figure 4). Being smaller in size, it is difficult to see the finer retinotopic details in these areas, however, the predicted maps in these areas do appear to reflect the general organizational features present in the empirical data. Figure 4 shows the average empirical polar angle maps (left and right hemispheres) of the test dataset as well as the average predicted maps. On these, we have explicitly drawn isopolar angle lines (dashed white lines) at phase reversals based on the average empirical maps and then positioned the grid identically on the predicted maps as well as on the visual hierarchy parcels as defined by the Wang et al. (2015) atlas. Note that visual areas boundaries are delineated by reversals in polar angle representation, near the horizontal meridian (0⁰ - left hemisphere; 180⁰ - right hemisphere), upper vertical meridian (90⁰) and lower vertical meridian (270⁰). As can be seen in Figure 4, our model was able to capture polar angle reversals beyond early visual cortex, as seen for IPS0 and IPS1 boundaries in the parietal portion of the visual cortex, hV4 ventrally, and TO2 in the lateral visual cortex. Notably, despite being unclear in the average empirical polar angle map, the polar angle reversal between IPS1 and IPS2 can be seen clearly in the average predicted map (solid black line, left hemisphere) further demonstrating that our model is able to predict legitimate retinotopic details beyond early visual cortex. Note, however, that this difference could also be simply due to our model learning the mean retinotopic organization between IPS1 and IPS2 from the training dataset rather than reflecting an accurate prediction at the individual subject level. Additionally, the polar angle reversal between V3b and LO1 seen in the average empirical map of the left hemisphere was absent in the average predicted map, suggesting that our model failed to capture this boundary.

**Figure 4.**
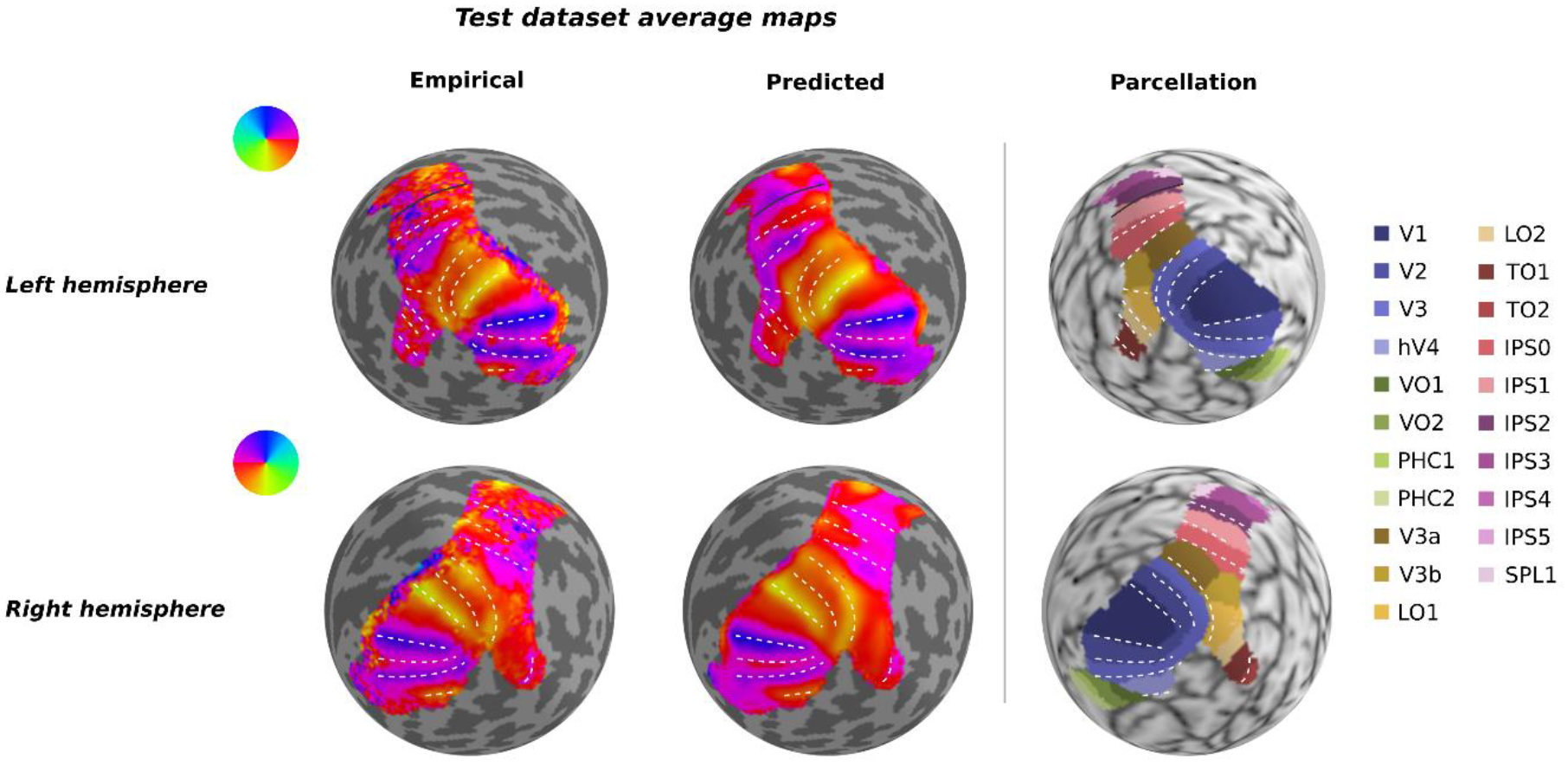
Polar angle reversals throughout the visual hierarchy. Upper row shows the average empirical polar angle map from the test dataset, the average predicted map, and the visual hierarchy parcels for the left hemisphere (adapted from Wang et al., 2015; see section 4.3 in the Methods). Lower row shows the same maps for the right hemisphere. The dashed white grid illustrates isopolar angle lines associated with boundaries of visual areas that were manually drawn based on the average empirical maps. The solid black line indicates an extra polar angle reversal between IPS1 and IPS2 seen in the average predicted map of the left hemisphere.

### 2.2 Individual variability in predicted maps

Our previous results demonstrate that our geometric deep learning models can predict the basic retinotopic organization of human visual cortex and that the predicted maps appear to be in line with the empirically measured maps for one participant, but are these models capable of predicting individual variability across participants? Figure 5 shows the results for 4 additional participants from the test dataset. For brevity, we only show the polar angle maps from the left hemisphere, but all the predicted maps for each test dataset participant are included as supplementary material (Supplementary Figures 1 and 2). In Figure 5a, the empirical polar angle maps are marked by unusual and/or discontinuous polar angle reversals. In the first three maps (Figure 5a, Participants 2-4), a discontinuous representation of the lower vertical meridian (that is, the boundaries between V1 and dorsal V2 as well as at the anterior border of dorsal V3) can be observed – indicated by the gray lines. Importantly, these unique variations were correctly predicted by our model. Perhaps even more striking, we see that these borders have merged to form a Y-shape for Participant 5, which was also observed in the predicted map. Importantly, a simple average map is unable to capture these clear examples of individual variability (Figure 5a, bottom row).

**Figure 5.**
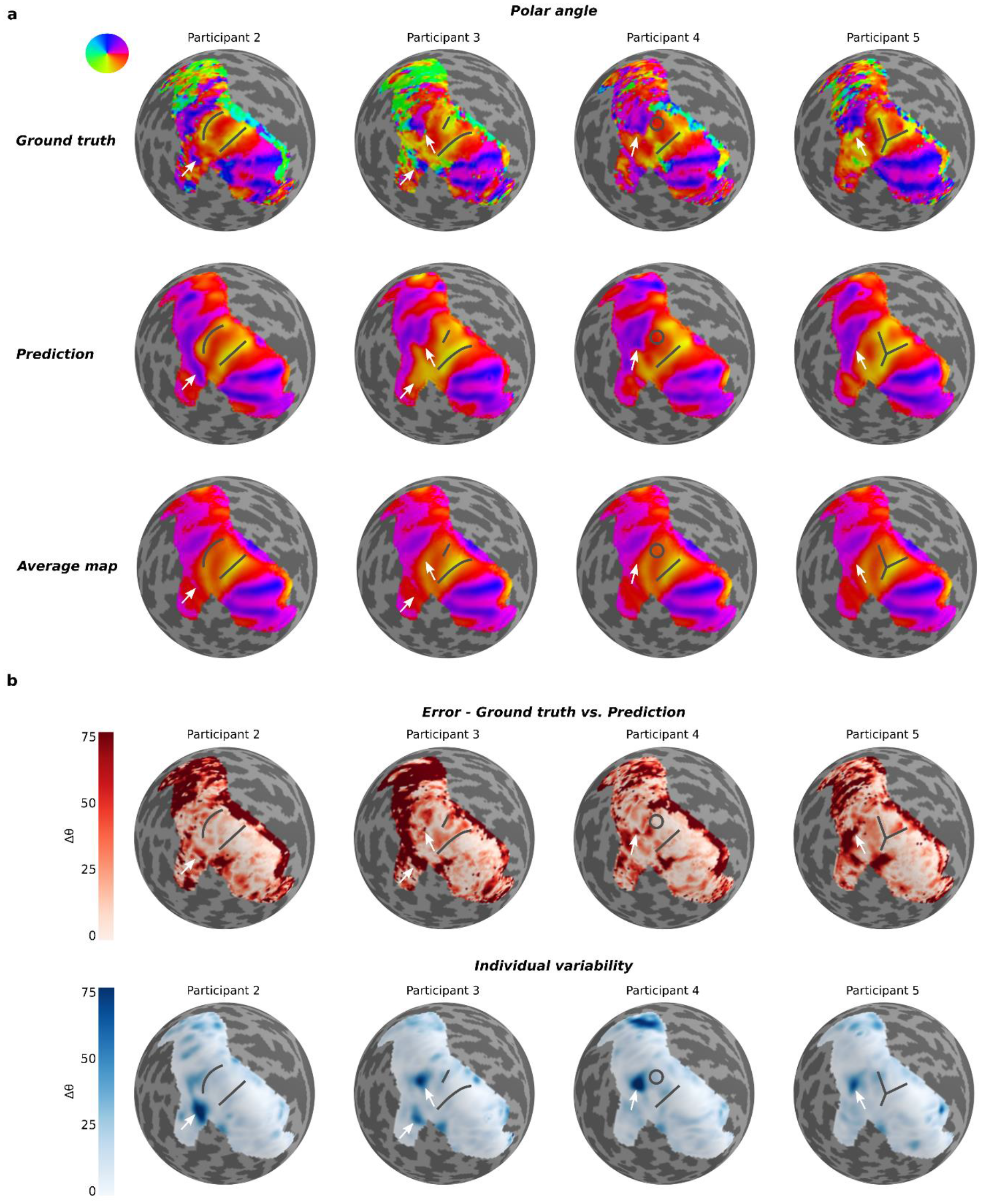
Individual variability in retinotopic maps. **a,** First row shows empirical (ground truth) polar angle maps and second row shows their respective predicted polar angle maps for four participants in the test dataset. The third row shows the average polar angle map from the training dataset. Grey lines indicate unique polar angle representations that were predicted by our model. **b**, Upper row shows the prediction error of each individual map and lower row shows how variable the same given map is from all the other predicted maps in the test dataset. White arrows indicate regions that differed the most among predicted maps.

To further assess the performance of our models, the difference (given by the smallest difference between two angles – Equation 2) between the predicted and the empirical (ground truth) polar angle values was determined in a vertex-wise manner (upper row, Figure 5b). This error estimate was then displayed on a standard cortical surface mesh to inspect where our models perform well and where they suffer. Predictions differed most from the ground truth near the foveal confluence (i.e., where the V1, V2, and V3 foveal representations merge), at the most peripheral eccentricities, and in some of the higher order visual areas (particularly those located dorsally in the intraparietal sulcus). Notably, each of these regions are known to be problematic when measuring retinotopic maps empirically. For example, it has been shown that retinotopic measurements within the foveal confluence are often not well represented due to instrumental and measurement limitations (Wandell and Winawer, 2010). The errors seen at the most peripheral eccentricities likely manifest from the suppressive effects seen in the cortex for retinotopic locations immediately outside the most peripheral limit of the visual stimuli (Puckett et al., 2014; Shmuel et al., 2002). Finally, the higher order areas are also known to be more difficult to map empirically using conventional procedures due to their smaller size and their higher degree of selectivity to specific types of stimuli (Wandell and Winawer, 2010). This brings us to an important caveat: the empirical maps, although treated as ground truth, may themselves misrepresent the underlying retinotopic organization. Accordingly, Figure 6 shows the average explained variance (R^2^) associated with the original pRF analysis of the empirical retinotopic mapping data (Benson et al., 2018) and as such, can be seen as a proxy for the quality of our ground truth. Pertinently, it shows that quality of the empirical data is indeed lowest in the regions where our model predictions were most error-prone – e.g., in the foveal confluence (Figure 6, arrow a), at the most peripheral eccentricities (Figure 6, arrow b), and in some of the higher-order areas (Figure 6, arrow c). Despite being a reasonable estimate of the quality of the data, the explained variance is an indirect and somewhat noisy measure. For example, the explained variance can be systematically high even when the pRF estimate is wrong – depending on details such as vessel location. Because the explained variance is systematically high in some parts of the visual cortex and systematically low in others, it can essentially be considered another visual field map. Here we use these maps to weight our loss function (see Methods); however, it would also be possible to train the network to predict these maps in addition to the eccentricity and polar angle maps.

**Figure 6.**
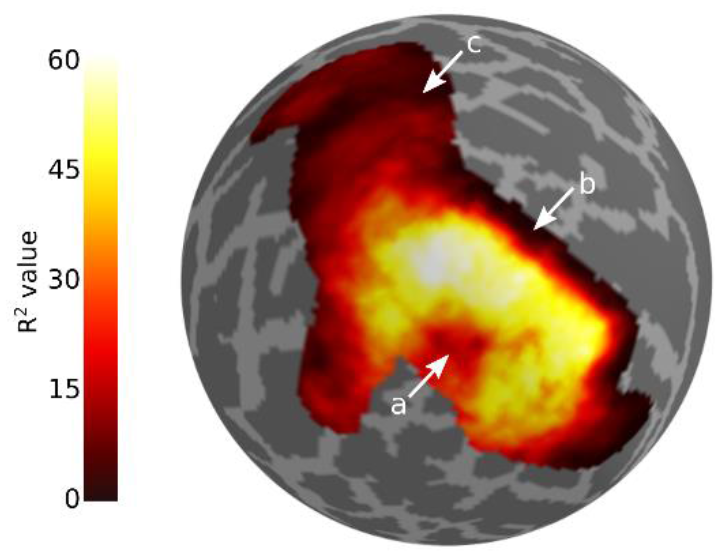
Quality of the empirical retinotopic mapping data. Average explained variance (R^2^) map of pRF modeling (Benson et al., 2018) from all participants in the test dataset. Arrows indicate the foveal confluence (a), peripheral eccentricities (b), and some of the higher-order areas (c) where explained variance is lowest.

Similar to the error estimates, we were interested in quantifying the extent to which each individual’s predicted map within the test dataset varied from the others. For this we computed an individual variability estimate as the difference (Equation 2) between a particular predicted map and each of the other predicted maps in the test dataset in a vertex-wise manner and averaged across all comparisons. Results are shown in Figure 5b, lower row. As expected, locations that more prominently differed among predicted maps in early visual cortex were the areas in and near the unique discontinuities seen around the lower vertical meridian lines in the polar angle maps (Figure 5, white arrows).

To further assess the accuracy of our individual predictions, we compared the performance of our model to a simple average of the retinotopic maps (computed using the training dataset, see Supplementary Figure 3) in predicting polar angle, eccentricity, and pRF center location. Here, the group-average maps were used as predictors of retinotopic organization and, as such, these predictions did not vary across individuals. To this end, we quantified the mean error over vertices within the range of 1-8⁰ of eccentricity within a few regions of interest: (1) dorsal V1-3 (Wang et al., 2015), where we see most of the unique variation in the early visual cortex (Figure 5 and Supplementary Figure 1), (2) the entire early visual cortex, and (3) higher order visual areas (Figure 7). Figure 7a shows a diagram of the three different error metrics used to quantify the vertex-wise difference between predicted and empirical (1) polar angle, (2) eccentricity, and (3) pRF center location values. The pRF center location is determined by both the polar angle and eccentricity values for each vertex, and as such, the mislocalization of the pRF center is not independent but rather a composite of the other two error measures. Note that the difference (i.e., distance in the visual field) between the predicted and empirical pRF center location was then divided by the empirical eccentricity value from each participant to prevent errors at high eccentricity from dominating the error metric (Benson and Winawer, 2018).

**Figure 7.**
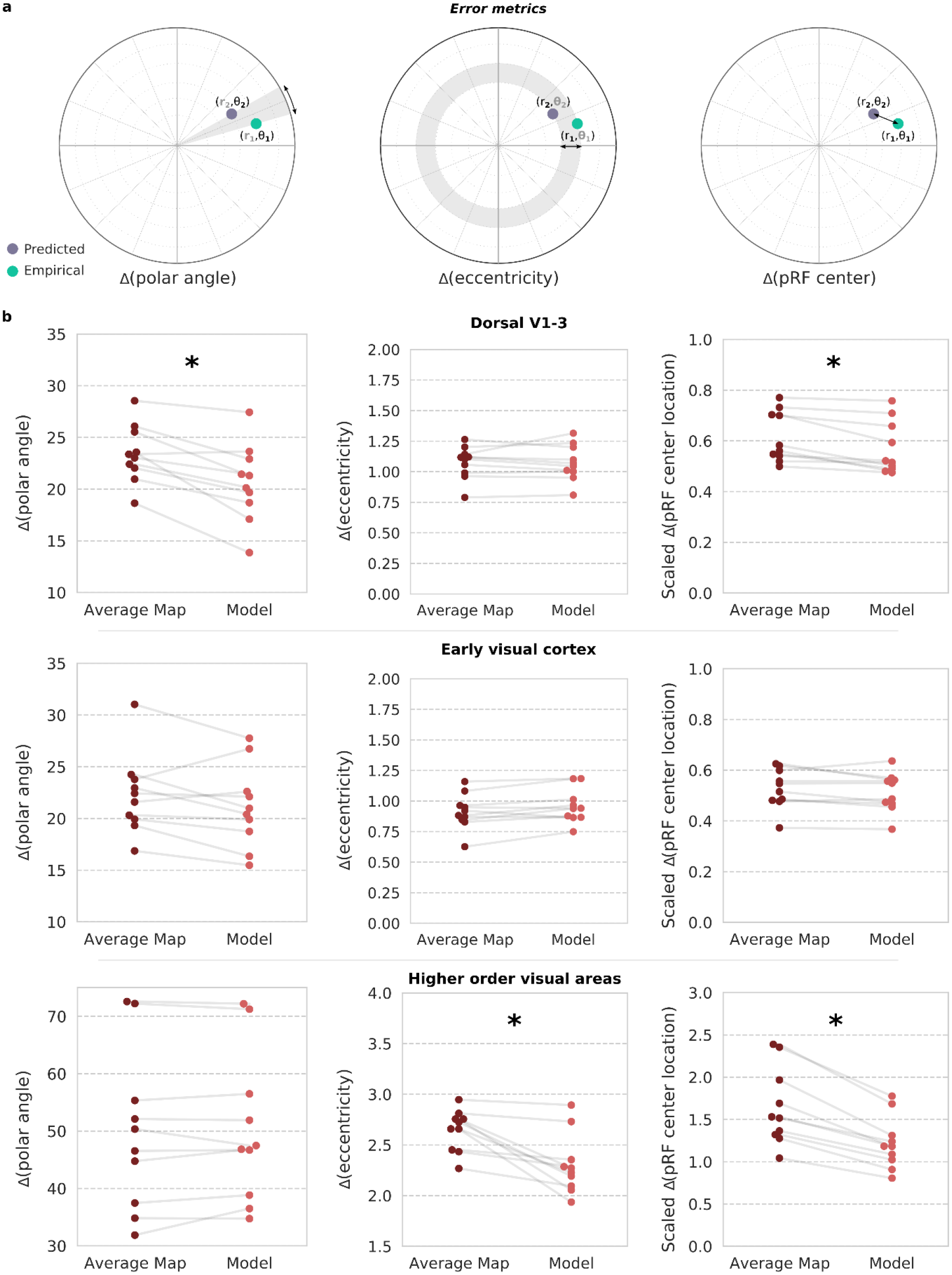
Prediction error based on predicting individuals’ retinotopic map using our model and using a group-average retinotopic map. **a,** Diagram of the three different error metrics used to quantify the vertex-wise difference between predicted (by either our model or an average map computed using the training dataset) and empirical polar angle (left), eccentricity (middle), and pRF center location (right) values. **b,** Vertex-wise errors were averaged over vertices within the range of 1-8⁰ of eccentricity within the dorsal portion of V1- 3 (top row), early visual cortex (middle row), and higher order visual areas (bottom row) for each individual from the test dataset (10 individuals). Dark red points illustrate the mean prediction error of group-average-based prediction for each individual, while light red points illustrate the mean prediction error of our models’ predicted maps. * indicates statistically significant differences between prediction errors from group-average and model-based predictions (i.e., p < 0.05).

The reduction in the mean error afforded by our model in the dorsal portion of V1-3 (Figure 7b) indicates that the variations generated by our model more accurately represent an individual’s polar angle map (p < 0.01, two-sided repeated measures t-test). However, this is less prominently seen at the entire early visual cortex (p = 0.11, two-sided repeated measures t-test), likely due the lower variability of the ventral portion of V1-3, and even less in higher order areas (p = 0.46, two-sided repeated measures t-test), where there is no clear improvement/deterioration in prediction accuracy with our model. Interestingly, while our approach did not generate more accurate eccentricity maps than a simple average map at the dorsal portion of V1-3 and the entire early visual cortex, we found a significant improvement with our model’s predictions at higher order visual areas (p < 0.01, two-sided repeated measures t-test). Finally, we found that using both the eccentricity and polar angle maps generated by our approach to estimate pRF center location was significantly more accurate than only using average maps in both the dorsal portion of V1-3 (p < 0.01, two-sided repeated measures t-test) and higher order visual areas (p < 0.001, two-sided repeated measures t-test). The mean prediction error averaged over the test dataset of the four predictive models in comparison to the average maps are summarized in Supplementary Table 1. The mean vertex-wise explained variance of the four predictive models are in Supplementary Table 2.

Finally, to further quantify the performance of our model in comparison to an average map and a well-established model of retinotopic organization (Benson et al., 2014), we determined the difference between polar angle predictions and different pRF fits (split halves; Table 1) for the left hemisphere at the dorsal portion of V1-3. Three distinct pRF fits were originally generated by Benson et al. (2018): one fit used the data from all six runs of the retinotopic mapping stimuli (i.e., the data used for our models’ training – fit 1); the second (fit 2) and the third fit (fit 3) used only the first and the second half of each of the six runs (i.e., split halves). We found that the mean difference between fit 1 and fit 2 and between fit 1 and fit 3 were 4.51 (SD = 1.06) and 5.50 (SD = 1.84), respectively. These differences are very low in comparison to the difference between predicted maps and empirical maps, which suggests that pRF estimates were consistent across fits. Therefore, we expected that predicted maps would perform equally well regardless of the empirical mapping data used to estimate the prediction errors.

**Table 1.** Mean prediction error of our model, the group-average map, and the Benson et al. (2014) template-based prediction across participants in the test dataset, while varying empirical mapping data using different population receptive field (pRF) fits (fit 1 = all data; fits 2 and 3 = split halves). Vertex-wise differences between predicted (our models’, a group-average map, or Benson et al. (2014) template map) and empirical polar angle (pRF fits 1, 2 and 3) maps were determined for the left hemisphere. Then, vertex-wise errors were averaged over vertices in the range of 1-8⁰ of eccentricity within the dorsal portion of V1-3, as in Wang et al. (2015), for each individual, and subsequently averaged across individuals in the test dataset (10 individuals). Note that retinotopic maps from pRF fit 1 were the actual data used for our model’s training.

| Prediction Type | Population receptive field modeling fit (mean $\Delta\theta \pm SD$) | | |
| --- | --- | --- | --- |
|  | Fit 1 | Fit 2 | Fit 3 |
| *Our model* | 20.62 ± 3.53 | 20.45 ± 3.79 | 21.18 ± 3.55 |
| *Average map* | 23.42 ± 2.64 | 23.62 ± 2.86 | 23.55 ± 2.53 |
| *Benson et al. (2014)* | 22.15 ± 4.57 | 21.79 ± 4.60 | 22.65 ± 4.26 |

Since our predictions were generated on a standard space (the HCP 32k fs_LR), all quantitative results were determined in the same space. After bringing Benson et al. (2014) model’s predictions to the standard space, predictions were nearly identical for all participants as this model warps template retinotopic maps (Supplementary Figure 4) to an individual’s native cortical surface. Therefore, after bringing these predictions back to a standard space, it is expected that the predicted maps will be nearly identical, with some minor differences due to the alignment procedure. We performed a 3 × 3 repeated measures ANOVA to test for effects of Prediction Type (our model, the group-average map, and the Benson et al. (2014) model) and Empirical Dataset used (pRF fit 1, fit 2, and fit 3). A main effect of Prediction Type was found (F(2, 18) = 5.222, p = 0.016; Supplementary Table 3), and the post-hoc test (using the Bonferroni correction to adjust p; Supplementary Table 4) indicated that our model’s predictions were significantly more accurate than the group-average-based predictions (p < 0.05). No statistically significant difference was found for the remaining comparisons.

### 2.3 Variability guided by anatomical features

To demonstrate the importance of the anatomical features in shaping individual differences in the functional organization of the visual cortex, we predicted three new sets of retinotopic maps using the previously trained model for each participant after manipulating the input by (1) rotating the anatomical feature maps, (2) randomly shuffling the anatomical input features, and (3) setting the input features equal to a constant value (equal to the mean curvature and myelin values) (see Figure 8a for examples of these manipulated feature maps). Generally, we expected that disrupting the input features in these ways would result in higher error across predicted maps. More specifically, (1) rotating the feature maps results in a set of statistically intact input features; however, the anatomical information is from outside of visual cortex and hence should not be predictive of the functional organization therein. (2) Random shuffling of the anatomical features produces inputs that retain the same set of values from within visual cortex; however, the statistical structure among this information will be destroyed. Hence, if the spatial disposition of anatomical features on the cortical surface is important for predicting individual variability in retinotopic maps, this too should lack predictive power compared to using intact anatomical features. Finally, (3) by completely removing the variability present in the input features it was expected that not only would prediction accuracy suffer due to the lack of intact, patterned anatomical features, but so would the model’s ability to predict any individual variability.

**Figure 8.**
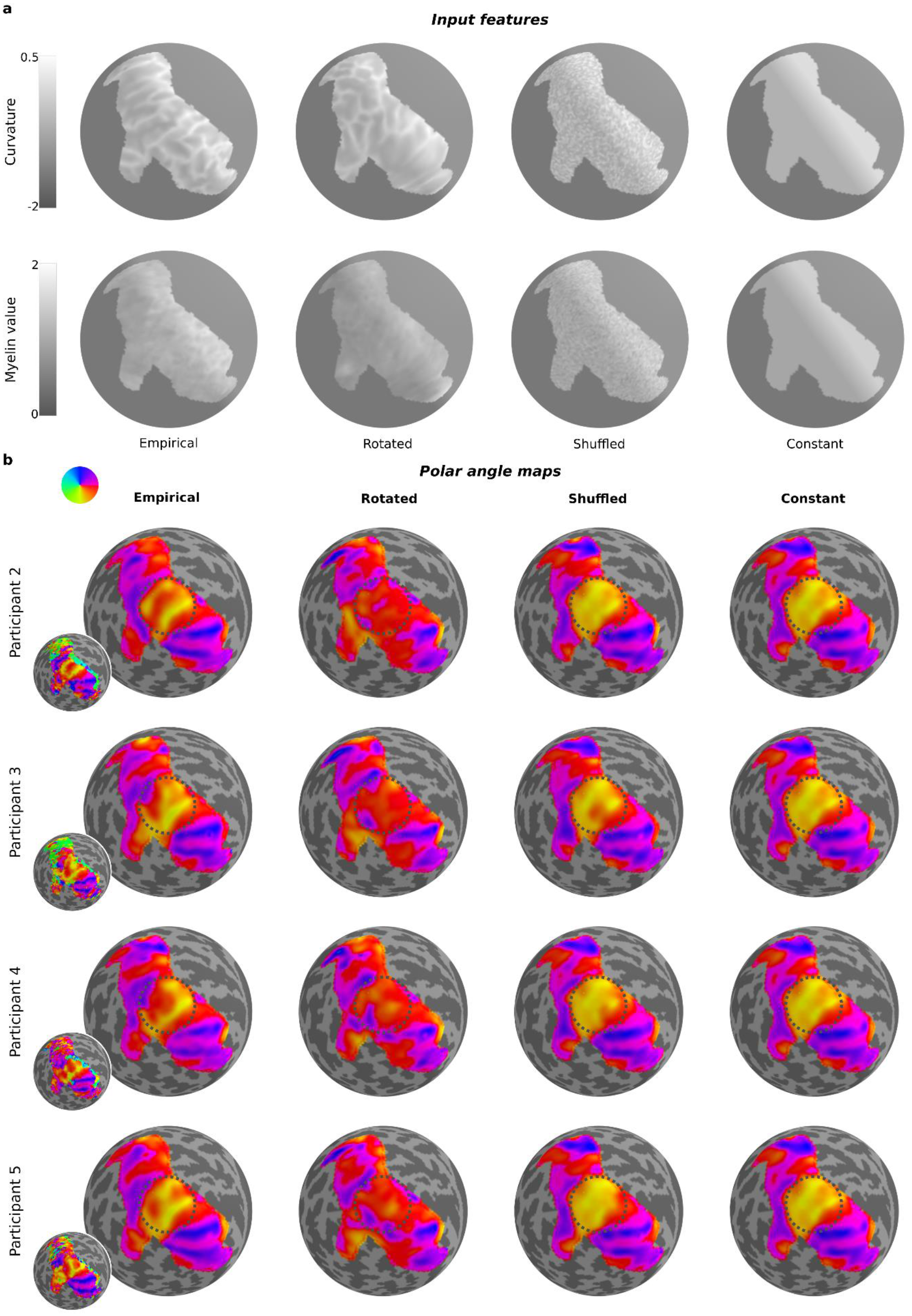
Individual variability in predicted maps is determined by the spatial arrangement and variability of anatomical features. **a,** An example of the new set of input features generated for each participant after (1) rotating the feature maps (second column), (2) randomly shuffling the anatomical input features (third column), and (3) setting the input features equal to a constant value (fourth column). **b,** Rows show the predicted polar angle maps using the empirical anatomical features (first column), and each of the three new sets of retinotopic maps generated using rotated feature maps (second column), shuffled features (third column), and input features equal to a constant value (fourth column) for participant 2-5. The inset images show empirical polar angle maps (ground truth). The dashed grey circle indicates the dorsal portion of early visual cortex. Overall, manipulating the input features generally leads to higher prediction errors (within individual) and lower individual variability (across individuals) in the new set of predicted maps than in those generated using intact input values, especially in the dorsal portion of the early visual cortex.

Contrasting the four sets of maps (Figure 8b) confirms the expectation that manipulating the input features generally leads to less accurate predictions compared to those generated using intact input values – especially in the dorsal portion of the early visual cortex where most of the variability was seen across individuals (Figure 5). As expected, using anatomical information from outside visual cortex (i.e., rotated input features) produced maps that are largely disrupted, with nearly no retinotopic structure matching the ground truth (Figure 8b – second column; see Supplementary Figure 5 for quantitative comparisons). Randomly shuffling anatomical features similarly led to less accurate predictions; however, the predicted maps are not nearly as disrupted as for the rotated input patterns (Figure 8b – third column). Without a structured input, it appears the model predictions tend toward the mean of the retinotopic organization learned by the network from the shape of the cortical surface. This likely reflects the fact that shuffled input features resemble constant input feature maps with the addition of unstructured noise, such that input values vary around the mean without any additional coherent spatial organization. Indeed, this can be further appreciated by examining the predictions associated with the constant inputs, which led to maps similar to those generated with shuffled input features (Figure 8b – fourth column). Here, again in a case where the statistical structure of the input data is lacking and is in no way biologically plausible, the model predictions contain recognizable retinotopic structure that tends toward the mean. However, it is important to note that despite these predictions being relatively accurate in some regions, they completely lack the nuanced differences seen across participants when using the intact anatomical feature patterns (or any differences at all in the case of using a constant input). Together, these results suggest that our model is relying on the spatial pattern of anatomical features to tune the general retinotopic organization learned from the shape of the cortical surface (i.e., surface topology) to generate idiosyncratic predictions.

### 2.4 Individual variability and genetic dependency

Although the HCP 7T retinotopy dataset contained mapping data for 181 different participants, not all datasets were strictly independent due to the presence of a large number of twin pairs (50 pairs of monozygotic and 34 pairs of dizygotic twins). For the previous results, we used data from all the participants to maximize the number of training examples seen by our network. However, given the genetic dependency among participants in our training and test data (all 10 of our test participants had a twin in the training set), we repeated the model training with a reduced training sample by excluding the twins of the participants in the test dataset from the training dataset. Importantly, the resulting model was still able to predict the retinotopic organization of visual cortex as well as unique variations in polar angle maps without exposure to the genetically-related mapping data (Supplementary Figure 6).

## 3 Discussion

This study shows that geometric deep learning can be used to predict the functional organization of visual cortex from anatomical features alone by building neural networks able to predict retinotopic maps throughout the visual cortical hierarchy. Remarkably the networks were able to predict unique features in these maps, showing sensitivity to individual differences. Practically, these predictions can be used to demarcate cortical visual areas within individuals according to known retinotopic mapping principles (Amano et al., 2009; Arcaro et al., 2009; DeYoe et al., 1996; Engel et al., 1997; Hansen et al., 2007; Sereno et al., 1995, 2001; Silver and Kastner, 2009; Swisher et al., 2007; Wandell et al., 2007; Wandell and Winawer, 2010). However, it is important to note that the utility of our current models extends well beyond finding area boundaries as the models were not trained to simply predict visual areas but to predict detailed, vertex-wise retinotopic maps across the visual areas. Hence, our predictions can also be used to examine cortical responses as a function of eccentricity and polar angle – all without collecting additional functional data. Pertinently, our pre-trained models and code are both available for those interested in applying our approach for their own purposes (see *Data and code availability statement* section for more).

### 3.1 Individual variability in retinotopic maps

Visual cortex has long been known to be organized retinotopically, and understanding the nature of the relationship between the spatial pattern on the retina and its cortical representation has been an active area of research. It has been shown that this relationship can be described using relatively simple models, at least for early visual cortex (Balasubramanian et al., 2002; Schira et al., 2010; Schwartz, 1977). It was then shown that these analytic models can serve as templates to be subsequently warped to an individual’s cortical anatomy to predict the functional organization of early visual cortex (Benson et al., 2014, 2012). Similar to our approach, these predictions are based on individual’s anatomical structure. However, this previous approach assumes a rigid structure-function relationship through the use of the template, and hence despite being able to account for gross structural differences across participants, it is unable to account for any idiosyncratic differences in the structure-function relationship. The dependency on the template also prevents the application of this approach to areas outside early visual cortex where mathematical models of the retinotopic organization are lacking. Following this work, Benson and Winawer (2018) built a Bayesian model for predicting retinotopic maps that addressed these shortcomings through the addition of a small amount of empirical mapping data (Benson and Winawer, 2018). That is, the interpolated template map is used as a prior that can be further adjusted based on collected functional data. Critically, while this approach is able to predict some of the idiosyncratic differences seen across individuals, it is unable to do so from anatomical information alone. See Supplementary Table 5 for a further comparison of our approach to these others.

Differently from the current state-of-the-art models of retinotopic maps, our model was capable of capturing the intricate structure-function relationship of the visual cortex without enforcing a spatially consistent mapping (i.e., a template). It is often assumed that the dorsal third tier visual cortex is occupied by a single continuous area (V3) that extends all the way along the dorsal border of V2 (Benson et al., 2014; Benson and Winawer, 2018; Wang et al., 2015). However, research suggests that there may be a more complicated organization of the dorsal third tier visual cortex in non-human primates (Angelucci and Rosa, 2015; Zhu and Vanduffel, 2019) as well as in humans (Arcaro et al., 2015). In fact, the same Y-shaped (or forked) lower vertical meridian that our model was able to predict in dorsal visual cortex of Participant 5 (Figure 5 and 8) has been reported by other researchers and in multiple individuals (Van Essen and Glasser, 2018). Electrophysiological recordings from a multiunit electrode study in macaques suggest that the dorsal third tier visual cortex is not comprised of a single continuous area, but rather two visual areas, with a discontinuity at the lower vertical meridian representation at the anterior border of dorsal V3 (Gattass et al., 1988). Although the organization of the dorsal third tier visual cortex remains to be further investigated in humans, the discontinuous representation of the anterior border of dorsal V3 has been seen in some individuals in published work from different labs (Allen et al., 2021; Arcaro and Kastner, 2015; Benson and Winawer, 2018; Van Essen and Glasser, 2018). Pertinently, even the more advanced Bayesian model (Benson and Winawer, 2018) is unable to predict these peculiarities in the dorsal maps – despite the addition of functional data. As reported by the authors, this inability stems from fact that the Bayesian model assumes a prior probability of zero to any map that differs topologically from the template (see Supplementary Figure 4). The unprecedented ability of our model to predict these discontinuous representations of the lower vertical meridian at the anterior border of dorsal V3 not only demonstrates the more flexible nature of our approach, but it also suggests that this discontinuity is likely a real structure-related variation, worthy of further investigation.

Finally, to show the importance of the anatomical features in shaping individual differences in the functional organization of the visual cortex, we investigated how different manipulation strategies of the input features would impact our model’s predictions (Figure 8). We found that spatially structured input features (i.e. intact anatomical information) from outside visual cortex (rotated input features) produced maps that were largely disrupted, with nearly no retinotopic structure matching the ground truth. On the other hand, spatially unstructured input features (randomly shuffled or constant features) also led to less accurate predictions, although predicted maps were not nearly as disrupted as for the rotated input patterns. These results suggest that our model is relying on the spatial pattern of anatomical features to tune the general retinotopic organization learned from the shape of the cortical surface (i.e., surface topology) to generate accurate predictions at the individual level. In other words, the ventral V1/V2 boundary, for instance, is highly conserved in its relative position across individuals, which is determined by the anatomical features (i.e., curvature and myelination). This suggests that our model has learned that this boundary goes in a particular absolute position on the cortical surface space, which explains why it was intact even in maps generated with shuffled, constant, and to some extent rotated input features. For areas with higher individual variability (i.e. dorsal portion of V1-3), the spatial arrangement of anatomical features seems to more directly inform the positioning of polar angle reversals, explaining their absence in maps generated with shuffled, constant, and rotated input features. While these analyses were exploratory, our findings make provision for hypothesis-driven interpretability investigations.

### 3.2 Applying our model to other datasets

We trained our models using input data aligned and resampled to the HCP 32k fs_LR standard surface space (Glasser et al., 2013), allowing vertex correspondence across individuals. This alignment consists of aligning native cortical surfaces to the standard space, then interpolating an individual’s anatomical information onto corresponding vertices of a standard cortical surface. Here we used the same surface for all individuals (HCP 32k fs_LR midthickness standard surface space), with the individual’s anatomical data interpolated (vertex features). Hence, to apply our model to other datasets, the individual’s native cortical surface must be aligned to the HCP 32k fs_LR space (for example, by using the Connectome Workbench command wb_shortcuts-freesurfer-resample-prep) before resampling the data (curvature and myelin values) from the individual’s native surface space to the HCP space (wb_command-metric-resample). Then, after applying our pre-trained models (see Code and Data availability statement) to this resampled data, the predicted retinotopic maps can be taken back to the individual’s native space by the reverse process.

### 3.3 Limitations and Future Directions

Although our neural network was able to predict the main features of retinotopic maps as well as nuanced differences across individuals, there were regions where prediction accuracy suffered. The highest errors were seen near the foveal confluence, at the most peripheral eccentricities, and in some of the higher order visual areas. Amongst the possible reasons for this discrepancy, we have already mentioned one: the network’s ability to generate accurate predictions is determined largely by the quality of the training data. The quality of the training data is reflected to some extent in the explained variance associated with the original pRF analysis (Benson et al., 2018), which suggests that the retinotopic mapping values given by the pRF analysis are least reliable (i.e., lowest explained variance) in exactly the same regions where our model predictions were most error-prone (Figure 6). Considering the intrinsic difficulty of mapping these regions and the fact that fMRI data cannot offer perfect representations of the underlying neuronal activity, we weighted the loss function by the fraction of explained variance from the pRF solution of each vertex (Benson et al., 2018; Benson and Winawer, 2018) (see Methods), and hence by design, our model was trained to generate better predictions in regions with more reliable pRF estimates. Despite the issues with the empirical mapping data in higher order visual areas, it is worth noting that our models still generated more accurate predictions of pRF center location than the group-average maps in these areas (Figure 7). This is particularly promising since, to the best of our knowledge, none of the methods available for predicting retinotopic organization beyond early visual cortex is more accurate than group-average maps. Future work using stimuli tailored for the activation of these regions could be performed to test the assertion that the predictions would improve by providing the neural network with higher quality training data.

Another possible reason for less accurate predictions in some regions of visual cortex, especially the higher-order areas, is that the functional organization of these regions may simply not be as strongly coupled with the underlying anatomical properties. The location and organization of many visual areas outside early visual cortex (V1/V2/V3) remain controversial, and this can be attributed to a variety of reasons, ranging from instrumental and measurement limitations (Wandell and Winawer, 2010) to data pre-processing steps (e.g., inaccurate cortical surface alignment). Hence, it has proven difficult to study the particular structure-function relationship in these areas compared to early visual cortex. A systematic assessment of the effects of training data on predictions may provide insight into the degree to which the functional organization is coupled to the underlying anatomical properties. For example, if model accuracy does not improve in the higher-order areas despite better training data, it may indicate that the functional organization is less tightly coupled with the underling anatomical properties – at least the ones used here (i.e., curvature and myelin). Note that in such case, and even for the regions that are well-predicted, another future direction to improve the model predictions might be to include different input features. Although we tested two anatomical features known to be related to retinotopic organization, other anatomical parameters can be drawn from T1-weighted images, such as cortical thickness, which can easily be incorporated as input features to train new models. Additionally, it remains to be further investigated if using individual-specific surfaces (Gopinath et al., 2018) as opposed to interpolating individuals’ anatomical features onto a standard surface affords an increment in prediction accuracy of retinotopic maps. Moreover, one could also include non-anatomical features known to vary according to retinotopic organization such as functional connectivity measures (Griffis et al., 2017; Park et al., 2018) in order to improve prediction accuracy.

Besides reducing the amount of resources required to obtain high-quality and idiosyncratic retinotopic maps, there are many other potential uses for CNNs as implemented here. Being able to predict vertex-wise retinotopic maps that can subsequently be used to identify individual visual areas could enable to infer the visual area parcellations directly by training the network to classify each vertex with respect to its known visual area rather than visual field location. Pertinently, the HCP dataset has recently been annotated with individual-specific boundaries for V1, V2, and V3 (Benson et al., 2021), which could be integrated directly into our geometric deep learning framework for this purpose. There are also a number of disorders with known correlates in their visual field map properties/activity patterns across the visual areas (e.g., prosopagnosia and dyslexia) (Demb et al., 1997; Witthoft et al., 2016). Accordingly, the learning objective of our model could be altered to predict disease state from appropriate data - e.g., by classifying developmental prosopagnosia and typical adults from pRF size maps (Witthoft et al., 2016). And finally, although we have primarily been discussing visual cortex, this approach can easily be deployed to achieve similar goals in other topographically organized areas in the brain, including somatosensory (Puckett et al., 2020; Sanchez Panchuelo et al., 2018) and auditory cortices (Da Costa et al., 2015; Saenz and Langers, 2014).

### 3.4 Geometric deep learning in neuroscience

A key aspect of this work was the use of a geometric approach, which permitted deep learning to be performed directly on the cortical surface model. Doing so, we were able to take advantage of the actual structure of the cerebral cortex and hence more appropriately represent the data to the neural network. Geometric deep learning provides tools that facilitate the filtering of features along the surface representation. Importantly, such an approach avoids the mixing of information across inappropriate spatial domains by allowing convolutions only along the cortical sheet. This stands in stark contrast to conventional deep learning approaches, which operate through convolution across Euclidean space. These Euclidean-based approaches are often used to extract information embedded in volumetric MRI data (i.e., a stack of 2D slices), which is particularly useful for brain segmentation tasks (Bontempi et al., 2020; Henschel et al., 2020). However, for the investigation of cerebral cortex function, the application of these models would result in the convolutions being performed such that the information of interest would be mixed across regions that are not spatially contiguous in the cortex (e.g., across sulci or even hemispheres), and across regions containing data that are irrelevant for the given task (e.g., white matter and cerebral spinal fluid). With this respect, it is important to note that it was not the goal of our study to demonstrate the superiority of geometric deep learning over more traditional forms of convolutional neural networks. Rather, it was our aim to demonstrate an apposite application of geometric deep learning in neuroscience (i.e., for a task associated with spatially patterned information distributed across the cortical surface).

Here, the geometric deep learning approach was used to model the relationship between structural brain data and functional brain data; however, the potential for this technique transcends this particular use case as a similar approach can be applied to model relationships among task-based activation patterns, functional connectivity, structural information, sensory spaces, behavioral measures, and much more. Importantly, it is not at all rare for data to better be represented in a non-Euclidean space than in the Euclidean one. For example, also within the field of neuroscience, brain functional networks are often represented as graphs (i.e., connectomes) in which connections between two brain regions do not necessarily represent the physical distance between them. Geometric deep learning, being also well-suited for graph data (Bronstein et al., 2017; Zhang and Bellec, 2019), offers the field a powerful tool given the growing interest in predicting aspects of human behavior from these functional connectivity graphs (Cai et al., 2020; Dadi et al., 2019; Finn et al., 2015; Fountain-Zaragoza et al., 2019; Gao et al., 2019; Shen et al., 2017).

## 4 Methods

### 4.1 Participant information and data overview

Comprehensive information about the HCP 7T retinotopy dataset is given elsewhere (Benson et al., 2018). Briefly, complete retinotopic mapping and structural data were acquired for 181 participants (109 females, age 22-35) following the HCP protocol (Benson et al., 2018; Van Essen et al., 2013). All participants had normal or corrected-to-normal visual acuity. Among these participants, 100 individuals are genetically confirmed monozygotic twins (50 pairs), 68 individuals are dizygotic twins (34 pairs), 4 individuals are non-twin siblings (2 pairs), and 9 individuals whose twin/siblings were not included and/or were not genetically tested. Structural image acquisition included T1w and T2w structural scans at 0.7 mm isotropic resolution at a customized Siemens 3T Connectome scanner (Van Essen et al., 2013). White and pial cortical surfaces were reconstructed from the structural scans using FreeSurfer (http://surfer.nmr.mgh.harvard.edu/) and were aligned across participants to the HCP 32k fs_LR standard surface space (Glasser et al., 2013). Myelin maps were determined by the ratio of T1w/T2w images (Glasser and Van Essen, 2011) and appropriately normalized to account for B1+ transmit effects (Glasser et al., 2013).

Whole-brain fMRI data were acquired using a Siemens 7T Magnetom scanner at a resolution of 1.6 mm isotropic and 1 s TR. The data were processed using the HCP pipeline (Glasser et al., 2013), which involved correction for head motion and EPI spatial distortion, alignment of the fMRI data with the HCP standard surface space, and denoising for spatially specific structured noise. The data produced (in CIFTI format) by the pipeline consists of 91,282 grayordinates: 32,492 cortical vertices per hemisphere and 26,298 subcortical voxels with approximately 2 mm spatial resolution. All data are publicly available on the ConnectomeDB database (https://db.humanconnectome.org).

### 4.2 Retinotopic mapping details

To build our deep learning model, we used previously analyzed retinotopic mapping data (Benson et al., 2018) – made available at https://balsa.wustl.edu/study/show/9Zkk as scene files. These files can be visualized using Connectome Workbench and are efficiently handled using the MATLAB GIFTI library and the Fieldtrip toolbox. For complete details on the experimental design and analysis procedure used to produce the maps, please refer to Benson et al. (2018). In brief, retinotopic stimuli were constrained to a circular region with a diameter of 16⁰. They consisted of slowly moving apertures of achromatic pink-noise background with dynamic colorful texture object, which was designed to elicit strong neural responses throughout the visual hierarchy – including in higher order visual areas. The experiment consisted of 6 runs with different set of apertures: wedges, rings and bars. Each aperture was presented twice, changing the direction of movement between runs, except for bar stimuli runs which were identical. Ring stimuli consisted of 8 cycles of a ring expanding away from the center or contracting towards the center with a period of 32s. Wedges stimuli consisted of 8 cycles of 90⁰ wedges rotating across the visual field counterclockwise or clockwise with a period of 32s. Finally, each bar stimuli run consisted of bars with different orientations (4 orientations) moving across different directions in the visual field. During each condition, participants were to attend to a semitransparent dot located at the center of the display and report whenever the color of the dot changed (via a button press) to ensure their gaze were fixed at the center of the display.

A pRF modeling procedure was then used to construct the retinotopic maps (Benson et al., 2018; Kay et al., 2013). Essentially, this modeling procedure estimates the spatial sensitivity profile within the visual field to which a grayordinate is responsive (i.e., its receptive field). For this, the fMRI time series elicited by the retinotopic mapping stimuli described above are modeled as the sum of a stimulus-related time series and a baseline time series. The stimulus-related response is then obtained by computing the dot product between the stimulus aperture time series and a 2D isotropic Gaussian (representing the pRF), applying a non-linearity component and rescaling the result by a gain factor, and then convolving it with a canonical hemodynamic response function. The modeled fMRI time series is optimized in order to best match the real fMRI time series by changing the 2D isotropic Gaussian parameters, such as its center position (x and y coordinates) and size, as well as the gain factor. Once the best parameters are determined, it is possible to construct retinotopic maps using each grayordinate’s preferred location in the visual field (i.e., its pRF’s center location). Polar angle maps reflect the polar angle in visual field to which a grayordinate is most responsive, while eccentricity maps reflect the distance from the center of the visual field (i.e., the fixation point) to which a grayordinate is most responsive. Note that the pRF modeling procedure is fully described in Benson et al. (2018).

### 4.3 Definition of the region of interest

As mentioned, the data files consisted of 91,282 grayordinates: 32,492 cortical surface vertices per hemisphere and 26,298 subcortical voxels. Here, we restricted the analysis to the cortical surface, hence ignoring the subcortical voxels. We further restricted our region of interest (ROI) to only include vertices within visual cortex using a surface-based probabilistic atlas (Wang et al., 2015). We modified the atlas slightly by extending the early visual cortex (V1, V2, V3) ROIs to include the foveal confluence (Schira et al., 2009) as well as combining ventral and dorsal components which are labelled separately in the original atlas (Figure 1). We also excluded the frontal eye fields (FEF) due to its discontinuity with the remaining areas. After selecting only vertices within this region of interest, we were left with 3,267 vertices for the left hemisphere (3,219 vertices of the right hemisphere), adding up to 19,024 connections (edges) between pairs of vertices (18,760 edges for the right hemisphere).

### 4.4 Data for deep learning

Our goal was to develop and train a deep neural network to predict the functional organization of the visual cortex (i.e., the retinotopic maps) from underlying anatomical properties (Figure 2). The anatomical data serving as input to the neural network included the mean curvature and myelin values (both of which have been shown to be related to the functional organization of visual cortex) resampled to the HCP 32k fs_LR standard surface space, which are explicit features. The connectivity among the ROI surface vertices (i.e., the surface topology) and their spatial disposition (i.e., their 3D coordinates) in the HCP 32k fs_LR standard surface space are also used by our model as implicit features (for more details, see Supplementary Methods). The output of the model was either the polar angle or eccentricity value for each vertex of the cortical surface model. Note that polar angle values are cyclical (i.e., 360⁰ = 0⁰). To avoid issues with the discontinuity in values, we shifted the polar angle values so that the left hemisphere was trained with the point of wrap-around (from 360⁰ to 0⁰) positioned at the horizontal meridian in the contralateral hemifield. After training, the values were shifted back to their original range.

For the modeling procedure, the data was represented on the midthickness cortical surface model (S1200_7T_Retinotopy181.L(R).midthickness_MSMAll.32k_fs_LR. surf.gii). Importantly, this model is a fiducial surface that maintains the actual geometry of the brain. Our results are, however, displayed using a spherical surface model (S1200_7T_Retinotopy181.L(R).sphere.32k_fs_LR.surf.gii) for visualization purposes. This permits all visual areas to be displayed in a single view.

### 4.5 Developing the neural network

Developing the neural network involved three main steps: (1) *training* the neural network, (2) hyperparameter *tuning*, and (3) *testing* the model. Prior to the training step, the 181 participants from the HCP dataset were randomly separated into three datasets: training (161 participants), development (10 participants) and test (10 participants) datasets (Supplementary Table 6). These datasets were used in each of the above three steps, respectively. During *training*, the network learned the correspondence between the retinotopic maps and the anatomical features by exposing the network to each example in the training dataset. Hence, the empirically-derived retinotopic maps served as the “ground truth” and the parameters of the neural network were optimized to minimize the difference between the predicted and empirical retinotopic maps. Model hyperparameters, such as the number of layers, were *tuned* (i.e., optimized) by inspecting model performance using the development dataset. Finally, once the final model was selected, the network was *tested* by assessing the predicted maps for each participant in the test dataset. This procedure was followed for each hemisphere and each type of retinotopic mapping data (i.e., polar angle and eccentricity) separately, resulting in four predictive models.

### 4.6 Model architecture and training

We implemented a spline-based convolutional neural network (SplineCNN) (Fey et al., 2018) on cortical surfaces using PyTorch Geometric (Fey and Lenssen, 2019), a geometric deep learning extension of PyTorch. Among the broad range of methods defining convolution operations on irregular structured data, such as graphs and surfaces, spline-based convolution is amidst the spatial filtering approaches. In these approaches, filters aggregate information locally, around a vertex’s neighborhood. These filters exploit information given by relative positions of the neighbor vertices with respect to the reference vertex in addition to the information encoded in the connectivity, edge weights, and vertex features. Thus, SplineCNN aggregates vertex features in local neighborhoods weighted by learnable parameters of a continuous kernel function (Fey et al., 2018).

Following the original study, we denote spline-based convolutional layers as SConv(*k*,M_in_,M_out_) of which *k* is the kernel size, M_in_ is the number of input feature maps, and M_out_ is the number of output feature maps (Figure 2). Our final model architecture included 12 convolutional layers with the number of feature maps increasing to a maximum of 32 feature maps per layer, and then decreasing back to one final feature map. We fixed the kernel size (*k*) to 25 for all layers, as predicted maps looked smoother than the ones generated with smaller kernel sizes. A non-linear activation function, the exponential linear unit (ELU), was used after each convolution layer. Batch normalization and dropout (p=0.10) were applied to all layers, except the last. Batch normalization is an approach commonly used in the field of machine learning to automatically normalize each layer’s input data, preventing the shift of the input data distribution during training, which results in faster training and lower variability among models (Frankle et al., 2020; Ioffe and Szegedy, 2015). Dropout refers to a regularization method used to prevent model overfitting (Srivastava et al., 2014). In our model, we only used Cartesian coordinates to compute relative distance between vertices and the degree of B-spline basis m = 1, due to the fact that models with those parameters showed better performance than the models using higher degree of B-spline basis or using spherical coordinates in previous experiments (Fey et al., 2018). Training was carried out for 200 epochs with a batch size of 1 and a learning rate at 0.01 for 100 epochs that was then adjusted to 0.005, using Adam optimizer. Our models’ learning objective was to reduce the difference between predicted retinotopic map and ground truth (i.e., the empirical retinotopic map). This mapping objective is measured by the smooth L1 loss function (Equation 1).

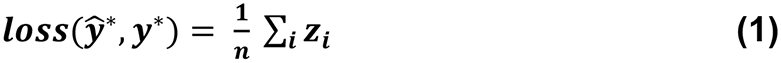

where *y*^∗^ and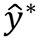 are, respectively, the empirical and predicted value weighted by individual-specific explained variance (R^2^) from the pRF modeling procedure (Benson et al., 2018), *n* is the total number of vertices, and *z_i_* is given by:

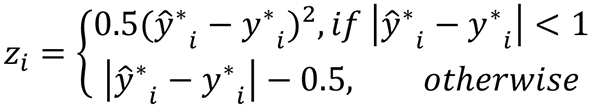

Models were implemented using Python 3.7.3 and Pytorch 1.2.0. Surface plots were generated using Nilearn, a Python module for fast statistical learning on neuroimaging data (Abraham et al., 2014). Training was performed on a high-performance computing cluster using NVIDIA Tesla V100 Accelerator units.

### 4.7 Hyperparameter tuning

Hyperparameter tuning was conducted by considering the performance of the model on predicting retinotopic maps of participants within the development dataset. We used the mean absolute error to compute the difference between predicted and expected values (ground truth). During the training stage, we monitored the decrease of the loss function (i.e., the prediction error on the training dataset) and the decrease of the mean absolute error (i.e., the prediction error on the development dataset). We also visually monitored predicted retinotopic maps using the development dataset, as we observed that despite the prediction error within the development dataset reaching a plateau around 100 epochs, some of the details in the predicted retinotopic maps continued to vary with additional training. To allow for some of this additional fine tuning, we fixed our training to 200 epochs.

To find the most appropriate number of layers, we trained five different models per number of layers, varying from 1 to 20 layers, as the performance of the final model fluctuated due to random initialization of the learnable parameters. We selected the number of layers based on the performance of the trained network, which was estimated by the smallest difference between two angles, given by:

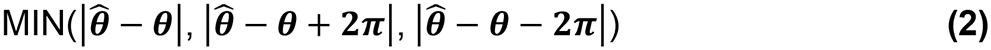

for 0< *θ* < 2*π*. To this end, we considered two factors: (1) how well model prediction matched the ground truth (error) and (2) how much a predicted map differed from the others (individual variability). To estimate the error, we computed the difference (given by Equation 2) between the predicted and the empirical (ground truth) polar angle values in a vertex-wise manner and averaged across all participants in the development dataset and within the various ROIs. The other factor, individual variability, was determined by the difference between a specific predicted map and each other predicted map in the development dataset in a vertex-wise manner and averaged across all combinations (there are 10 participants in the development dataset, hence 9 combinations), and then averaged across participants and within the various ROIs. This was performed for all models (5 models per number of layers) and we selected the number of layers that resulted in the best performance across all the ROIs in both metrics (12 layers; see Supplementary Figure 7). This selection was based on mapping polar angle values in the left hemisphere; however, we tested a range of layers using the right hemisphere data and found similar results. Finally, once the appropriate number of layers was established, we chose the best model among the 5 models that were trained with the same hyperparameters but with different learnable parameters (due to the random initialization) to evaluate on the test dataset. For the eccentricity models (left and right hemispheres), we kept the same hyperparameters as optimized for polar angle models.

### 4.8 Model testing and evaluation metric

The selected models were evaluated based on their performance on the test dataset considering the same two factors, error and individual variability. Importantly, the test dataset was never seen by the network nor by the researchers until the deep learning architecture was finalized and the final model selected. Again, the error was determined by the difference between the predicted and the empirical polar angle values in a vertex-wise manner and averaged across all participants – this time in the test dataset. Individual variability was determined by the difference between a specific predicted map and each other predicted map in the test dataset in a vertex-wise manner and averaged across all combinations (9 combinations), and then averaged across participants.

### 4.9 Model performance in comparison to an average map

To assess the accuracy of our individual predictions, we compared the performance of our model to a simple average of the retinotopic maps in predicting polar angle, eccentricity, and pRF center location. We quantified the mean error over vertices within the range of 1-8⁰ of eccentricity, which was selected using the group-average eccentricity map over the training dataset. This range of values was chosen because, in the original pRF mapping experiment of the HCP, the visual stimulus extended to 8° of eccentricity. Additionally, due to the inherent difficulty in mapping the foveal confluence (Wandell and Winawer, 2010), we constrained our comparison to eccentricity values above 1°. Polar angle and eccentricity prediction errors were determined as described previously. PRF center location prediction error was determined by the Euclidean distance between two points in polar coordinates, given by:

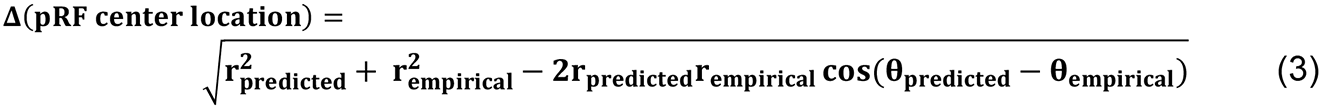

where r_predicted_ and θ_predicted_ are the predicted eccentricity and polar angle values, and r_empirical_ and θ_empirical_ are the empirical eccentricity and polar angle values. Then, the pRF center location errors were divided by the empirical eccentricity values from each participant to prevent errors at high eccentricity from dominating the error metric (Benson and Winawer, 2018).

After computing the vertex-wise prediction error, errors were averaged within a few regions of interest for each participant: (1) dorsal V1-3 - not including the foveal confluence (Wang et al., 2015), (2) the entire early visual cortex, and (3) higher order visual areas. The mean error across participants was determined to evaluate the performance of our model in comparison to a simple average map (Supplementary Table 1). We performed statistical comparisons with repeated measure t-test using Scipy (Virtanen et al., 2020). Additionally, we determined the vertex-wise explained variance of our model over the test dataset (Supplementary Table 2). First, individuals’ predicted and empirical maps were concatenated, which resulted in two matrices of size ‘number of vertices’ × 10. Then, we calculated the square of the correlation coefficient between each vertex’s (rows) predicted values and its corresponding empirical values.

Finally, we also compared our predictions with different pRF fits provided by the HCP. For each individual subject, Benson et al. (2018) performed three separate model fits: one fit used all data from six runs of retinotopic mapping stimuli (used here as our ground truth – fit 1), the second fit (fit 2) and the third fit (fit 3) used only the first half and only the second half of each of the six runs (i.e., split halves), respectively. Prediction error was determined between our model and pRF fits 1 to 3, between the average map and pRF fits 1 to 3, and between Benson et al. (2014) model and pRF fits 1 to 3. We report the mean vertex-wise prediction error within the dorsal portion of V1-3 averaged over the test dataset (Table 1). Statistical comparison was performed with two-way (Prediction Type X Empirical Dataset) repeated measures ANOVA using Jamovi (The jamovi project, 2021) (Supplementary Table 3 and 4).

### 4.10 Assessing spatial organization of the anatomical features

To evaluate the importance of the spatial organization of the anatomical features and their variability for the prediction of individual differences in retinotopic maps, we tested the effects of modifying the input data in three ways using the previously trained model: (1) rotating the feature maps, (2) randomly shuffling the anatomical features, and (3) setting the input features equal to a constant value. (1) To rotate the feature maps, AFNI’s ConvertSurface was used with the flag ‘-xmat_1D NegXY’ (which flips the sign of X and Y coordinates), followed by the SurfToSurf command that interpolates the rotated surface nodes onto the original surface. This procedure essentially replaces the intact curvature and myelin patterns in visual cortex with intact (but inappropriate) patterns from elsewhere in the brain. (2) Random shuffling of the input features, in contrast, uses the values from the intact feature maps in visual cortex; however, the values are permuted across the ROI vertices such that the spatial organization of the anatomical information was completely disrupted. (3) To test the effect of completely removing the variability present in the input features, all curvature and myelin values of each participant in the test dataset was set equal to the mean curvature (0.027) and the mean myelin (1.439) values, respectively. Note that the mean values were computed using all participants from the training dataset.

## Ethics statement

We analyzed structural and functional data recorded from 181 participants in the Human Connectome Project (http://www.humanconnectome.org) (Benson et al., 2018; Van Essen et al., 2013). Participant recruitment and data collection were carried out by Washington University and the University of Minnesota. The Institutional Review Board (IRB) at Washington University approved all experimental procedures (IRB number 201204036; “Mapping the Human Connectome: Structure, Function, and Heritability”), and all participants provided written informed consent before data collection (Van Essen et al., 2013).

## Data and code availability statement

The data used in this study is publicly available at BALSA (https://balsa.wustl.edu/study/show/9Zkk). All pre-trained models will be available on the Open Science Framework (https://osf.io/95w4y/) and all accompanying Python source code will be available upon publication on GitHub (https://github.com/Puckett-Lab/deepRetinotopy). Moreover, an executable code notebook hosted on Google Collab is available that allows anyone to load the trained models and to generate predictions on the test dataset (https://colab.research.google.com/drive/1zXrK5HK806iXkJ6IIKeg0ePKm1SCTQlB?usp=sharing). Note that the application of our model is not restricted to the HCP retinotopy dataset; however, it does require the input data (curvature and myelin values) to be resampled from an individual’s native space to the HCP standard space prior to computing the prediction.

## Credit authorship contribution statement

**Fernanda L. Ribeiro:** Conceptualization, Methodology, Software, Validation, Formal Analysis, Investigation, Data Curation, Writing – Original Draft, Review & Editing, Visualization. **Steffen Bollmann:** Conceptualization, Resources, Writing – Review & Editing. **Alexander M. Puckett:** Conceptualization, Methodology, Resources, Writing – Original Draft, Review & Editing, Supervision, Project administration, Funding Acquisition.

## Acknowledgments

This work was supported by the Australian Research Council (DE180100433). Data were provided by the Human Connectome Project, WU-Minn Consortium (Principal Investigators: David Van Essen and Kamil Ugurbil; 1U54MH091657) funded by the 16 NIH Institutes and Centers that support the NIH Blueprint for Neuroscience Research; and by the McDonnell Center for Systems Neuroscience at Washington University. The authors acknowledge Jake Carroll and Irek Porebski for their great support with the high-performance computing facilities of the Research Computing Centre at the University of Queensland.

