## Supplementary Material for "Predicting the retinotopic organization of human visual cortex from anatomy using geometric deep learning"

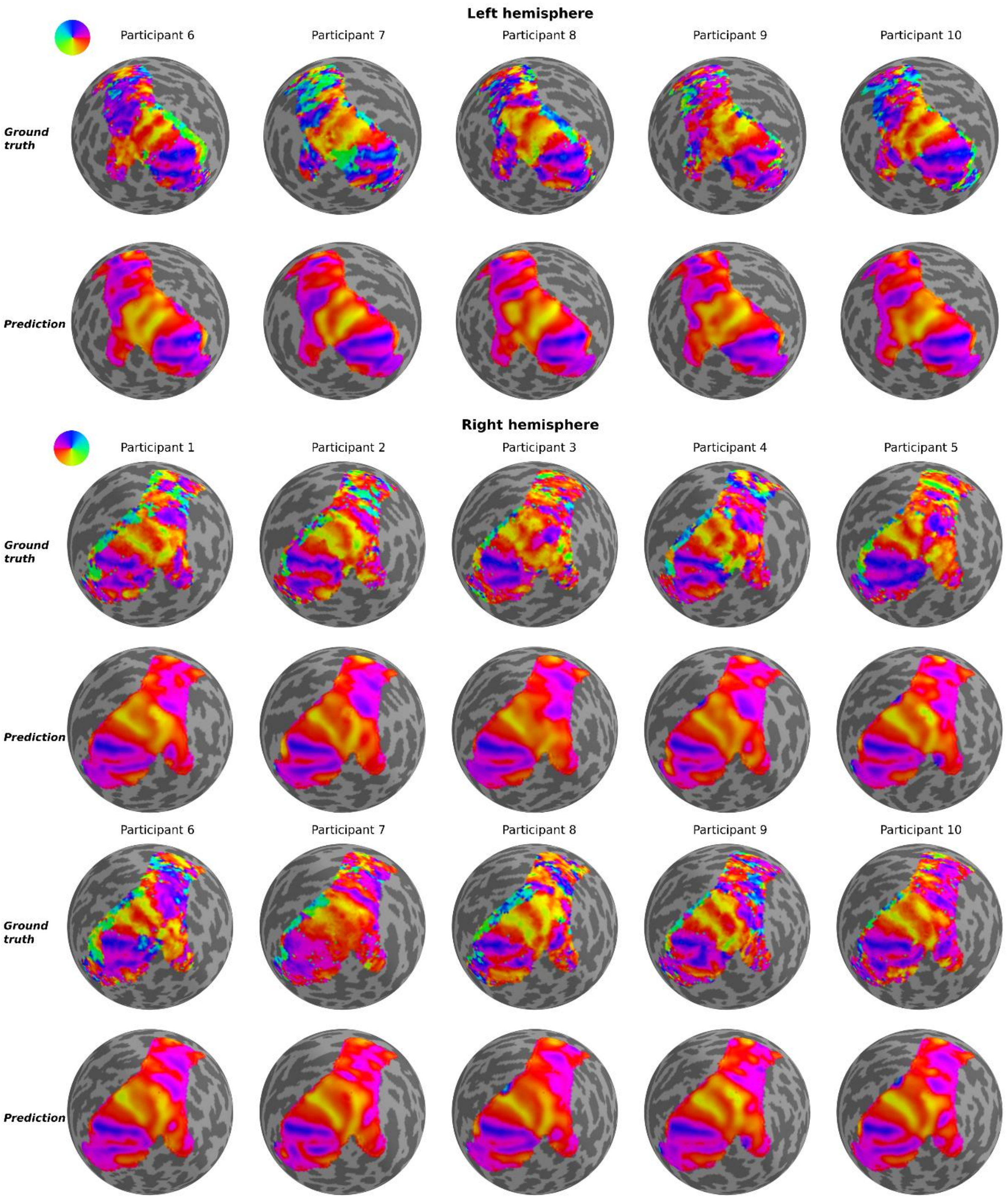


**Supplementary Figure 1 – Polar angle mapping with geometric deep learning.** Upper panel shows empirical (ground truth) and predicted polar angle maps for the left hemisphere of the remaining individuals in the test dataset (polar angle maps of participants 1-5 are in the main manuscript). Lower panel shows empirical and predicted polar angle maps for the right hemisphere of all individuals in the test dataset. Polar angles vary from 0º to 360º as indicated by the color wheels.

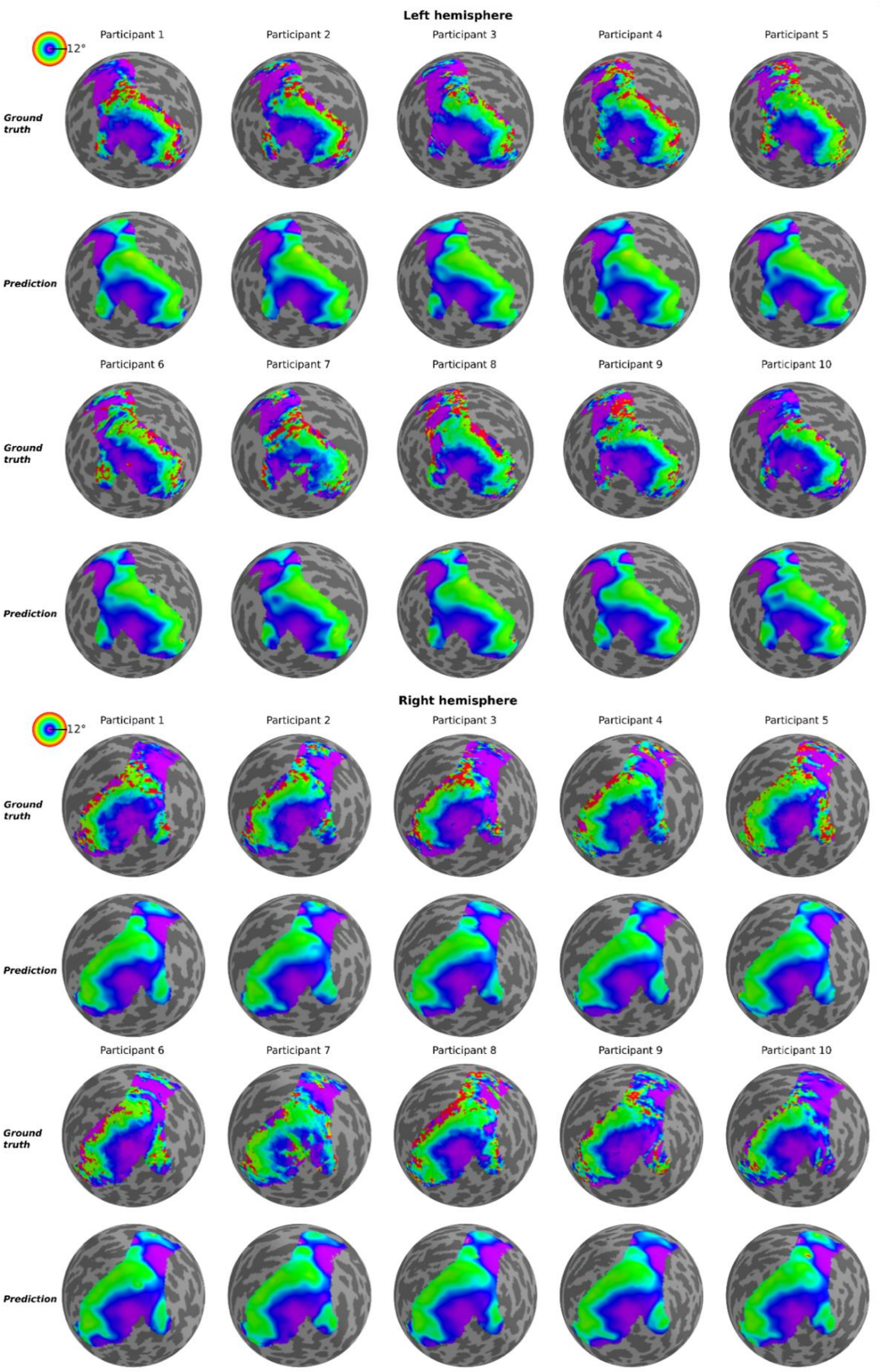


**Supplementary Figure 2 - Eccentricity mapping with geometric deep learning.** Upper panel shows empirical (ground truth) and predicted eccentricity maps for the left hemisphere of all individuals in the test dataset. Lower panel shows empirical and predicted polar angle maps for the right hemisphere of all individuals in the test dataset. Eccentricity varied from 0º to 12º as indicated by the color wheels.

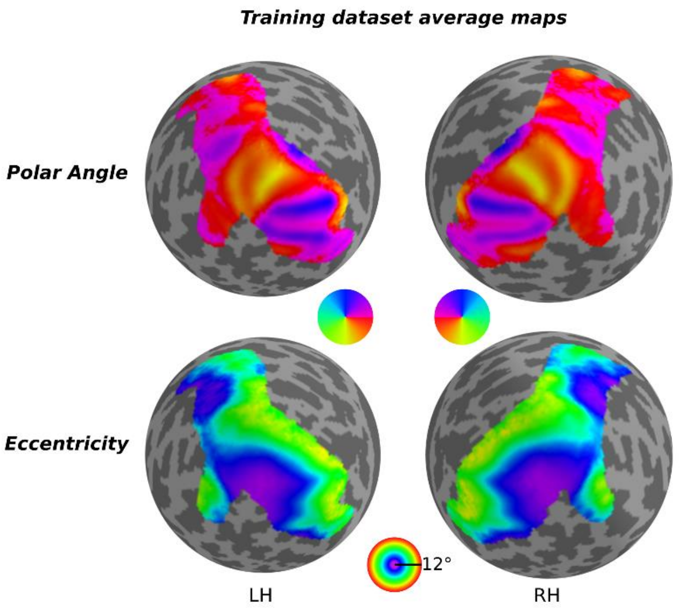


**Supplementary Figure 3 - Average retinotopic maps from training dataset.** Average polar angle and eccentricity maps from the training dataset used for comparisons with our models.

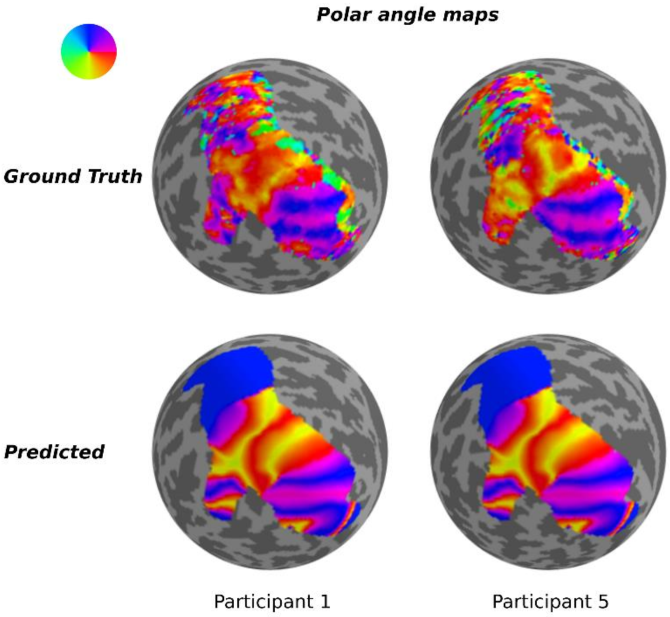


**Supplementary Figure 4 – Polar angle maps from two participants of the test dataset.** Upper row shows empirical (ground truth) polar angle maps from Participant 1 and Participant 5 of the test dataset. Bottom row shows predicted polar angle maps generated with Benson et al. (2014) model after resampling the data to the HCP 32k fs_LR template space. Note that all our predictions were generated on a template space (the HCP 32k fs_LR) and, as such, all the quantitative results were determined in the same template space. After bringing Benson et al. (2014) model's predictions to the template space, predictions were nearly identical for all participants, as expected.

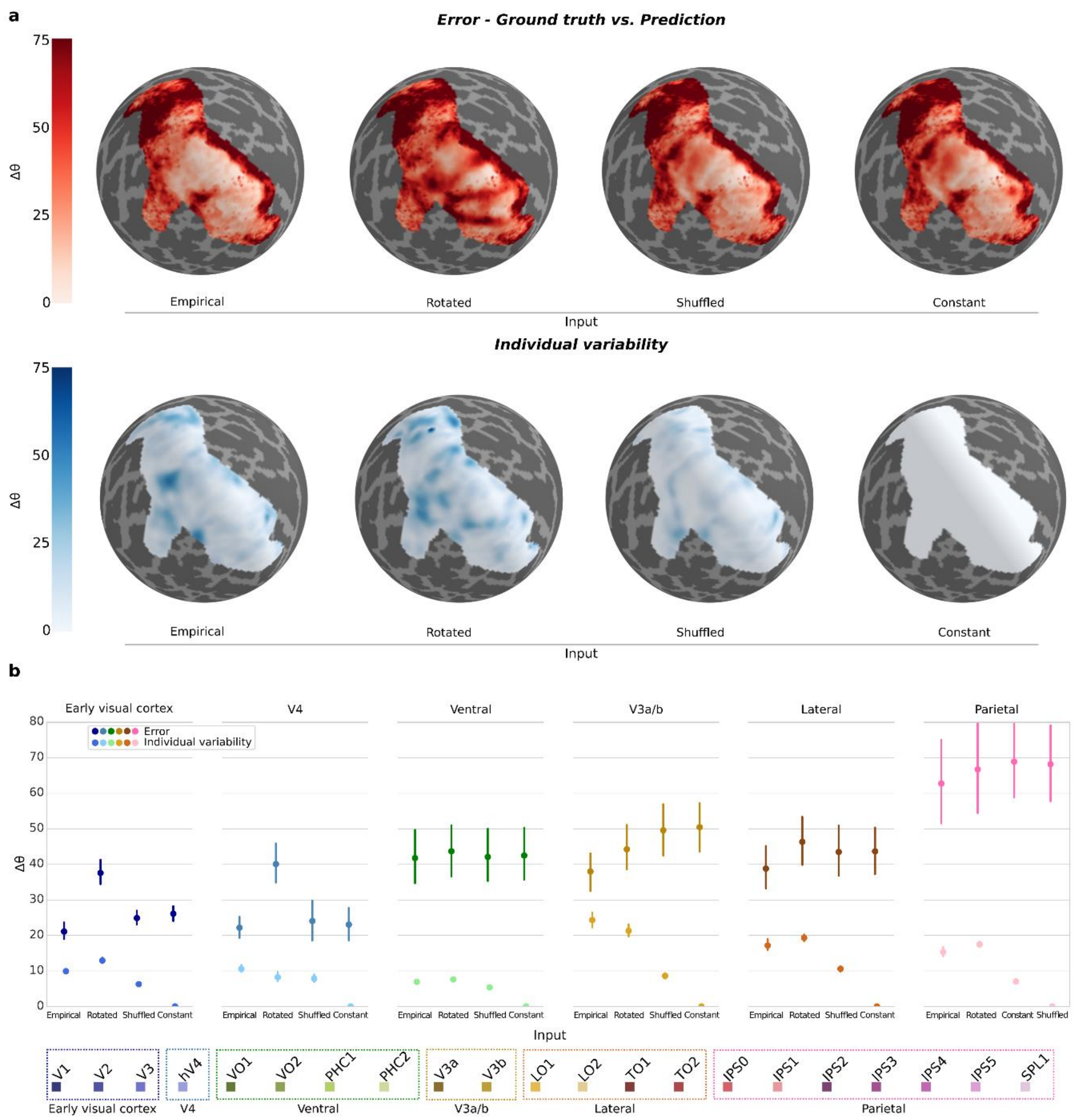


**Supplementary Figure 5 – Individual variability in predicted maps is determined by anatomical features. a,** Upper row shows the prediction error across participants in the test dataset using the empirical input features, rotated feature maps, shuffled input features, and constant input features. Lower row shows how variable predicted maps are from all the other predicted maps in the test dataset on average for each input condition. **b,** Mean and bootstrapped (n=1,000) 95% confidence interval of the average error and individual variability of predicted maps for each input condition within different visual area clusters.

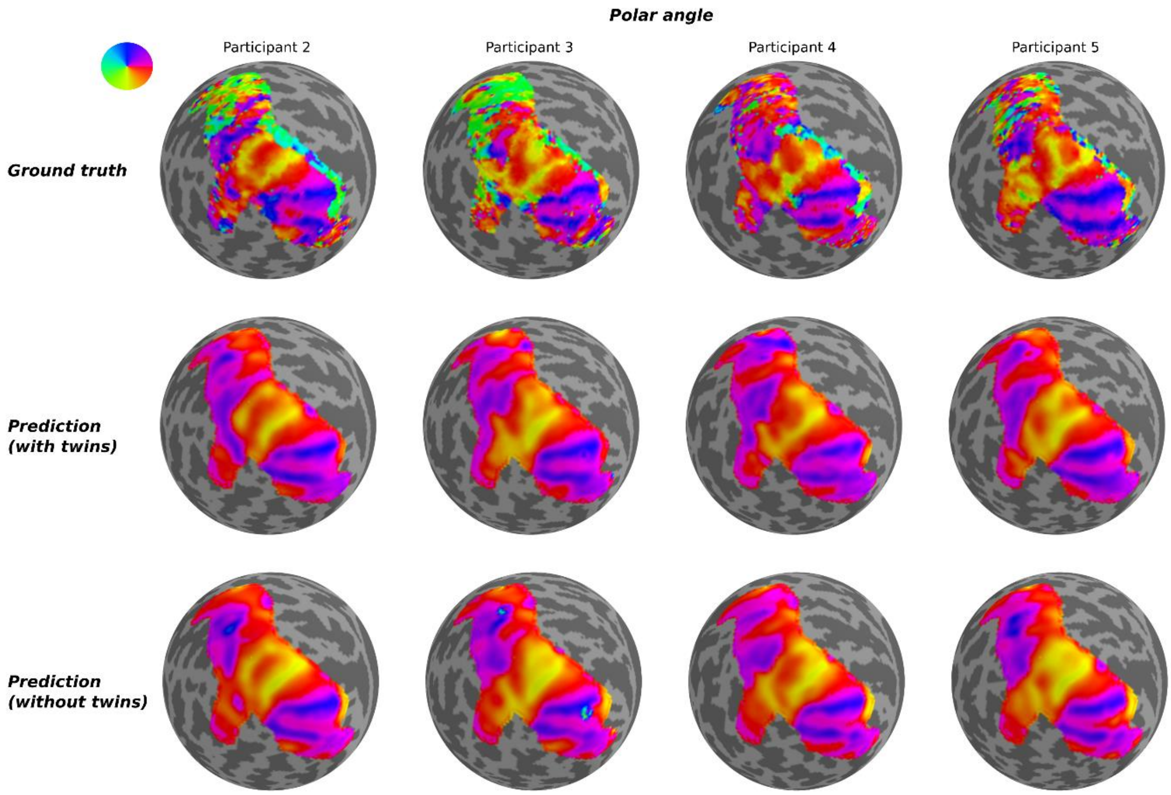


**Supplementary Figure 6 - Individual variability in retinotopic maps.** Upper row shows empirical (ground truth) polar angle maps for four participants in the test dataset, middle row shows their respective predicted polar angle maps generated by a model trained with their twins in the training dataset (Figure 4), and lower row shows predicted polar angle maps of the same participants generated by a model trained without their twins in the training dataset.

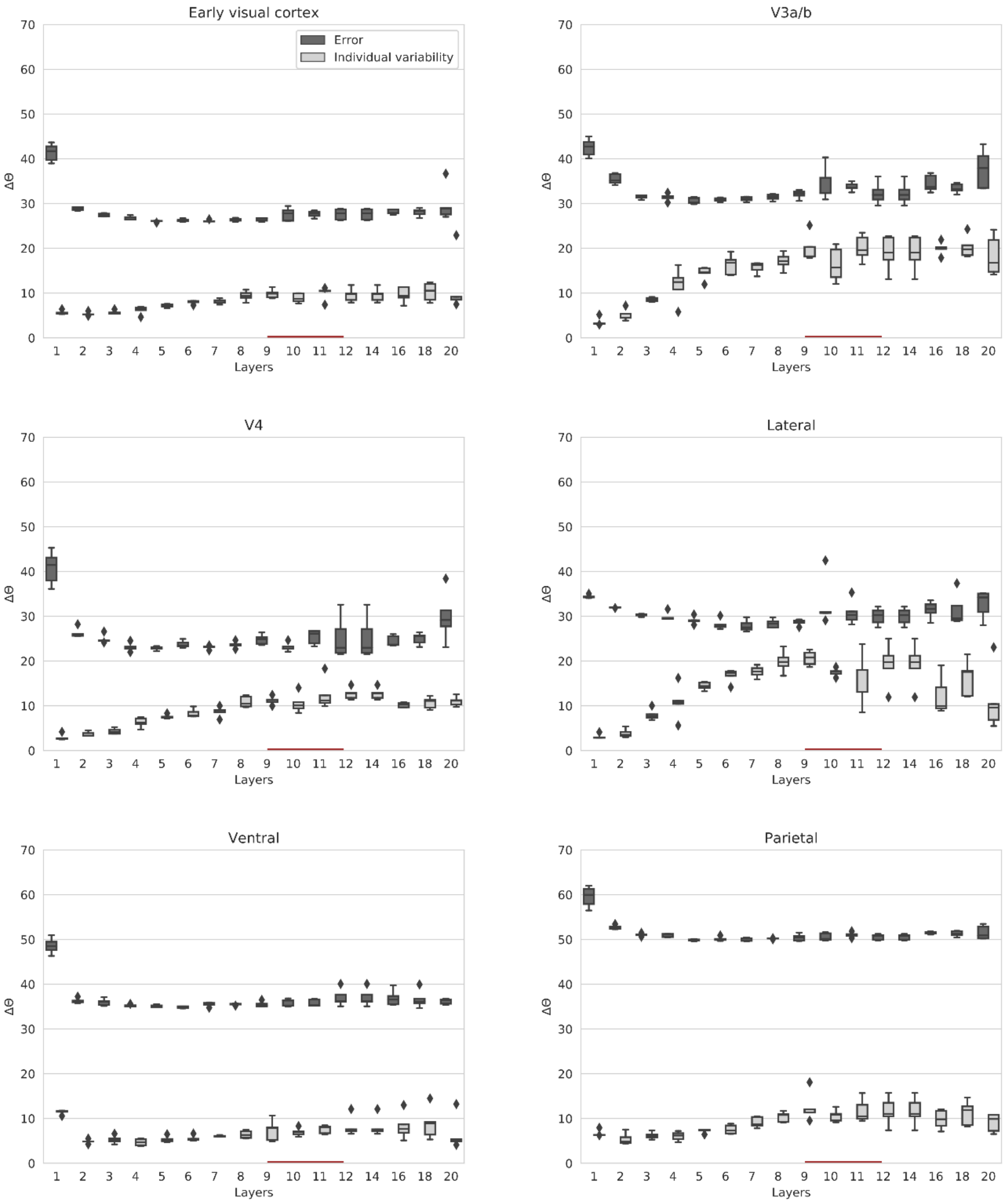


**Supplementary Figure 7 – Model performance with varying number of layers.** Box plots represent the mean prediction error (dark gray) and mean individual variability (light gray) of 5 different models trained for polar angle mapping in the left hemisphere – separated by visual area cluster. Evaluation of the performance of the models was conducted using the development dataset, as part of the hyperparameter tuning step. Models varied in the number of convolutional layers. Note that the mean error reduces as models get deeper (i.e., more layers) while the individual variability increases. However, performance starts to deteriorate for models with more than 12 layers. Models with 9 to 12 layers showed the best performance across all visual areas clusters.

**Supplementary Table 1 - Mean prediction error of the four predictive models versus the average maps on the test dataset.** Vertex-wise difference between predicted (by either our model or an average map) and empirical polar angle, eccentricity, and pRF center location values were determined for both left and right hemispheres. Then, vertex-wise errors were averaged over vertices within the range of 1-8º of eccentricity (as given by the average eccentricity map over the training dataset) within the dorsal portion of V1-3, early visual cortex, and higher order visual areas for each individual and subsequently averaged across individuals in the test dataset (10 individuals). Statistical significance was determined with two-sided repeated measures t-tests. Note: Alpha-level corrected for multiple comparisons per hemisphere is equal to 0.05/9=0.006.

| | | | | |
|---|---|---|---|---|
| | | **POLAR ANGLE** | | |
| | | **Mean Δθ ± std** | | **p-value** |
| *Hemisphere* | *Areas* | *Model* | *Average Map* | |
| LEFT | Dorsal V1-3 | 20.62 ± 3.53 | 23.42 ± 2.64 | 0.001 |
| | Early visual cortex | 21.10 ± 3.76 | 22.24 ± 3.62 | 0.111 |
| | Higher order areas | 50.28 ± 12.47 | 49.80 ± 13.39 | 0.464 |
| RIGHT | Dorsal V1-3 | 17.08 ± 3.03 | 19.13 ± 3.66 | 0.015 |
| | Early visual cortex | 19.38 ± 3.58 | 19.44 ± 2.77 | 0.880 |
| | Higher order areas | 39.15 ± 4.99 | 39.48 ± 5.70 | 0.387 |
| | | **ECCENTRICITY** | | |
| | | **Mean Δθ ± std** | | **p-value** |
| *Hemisphere* | *Areas* | *Model* | *Average Map* | |
| LEFT | Dorsal V1-3 | 1.07 ± 0.14 | 1.07 ± 0.13 | 0.945 |
| | Early visual cortex | 0.96 ± 0.13 | 0.91 ± 0.14 | 0.033 |
| | Higher order areas | 2.30 ± 0.28 | 2.65 ± 0.19 | 0.003 |
| RIGHT | Dorsal V1-3 | 0.83 ± 0.13 | 0.86 ± 0.18 | 0.509 |
| | Early visual cortex | 0.80 ± 0.11 | 0.80 ± 0.12 | 0.885 |
| | Higher order areas | 2.11 ± 0.30 | 2.36 ± 0.27 | 0.009 |
| | | **SCALED PRF CENTER LOCATION** | | |
| | | **Mean Δθ ± std** | | **p-value** |
| *Hemisphere* | *Areas* | *Model* | *Average Map* | |
| LEFT | Dorsal V1-3 | 0.57 ± 0.10 | 0.62 ± 0.09 | 0.003 |
| | Early visual cortex | 0.51 ± 0.07 | 0.53 ± 0.07 | 0.144 |
| | Higher order areas | 1.22 ± 0.29 | 1.65 ± 0.43 | <0.001 |
| RIGHT | Dorsal V1-3 | 0.48 ± 0.07 | 0.52 ± 0.09 | 0.015 |
| | Early visual cortex | 0.47 ± 0.06 | 0.49 ± 0.07 | 0.060 |
| | Higher order areas | 1.04 ± 0.20 | 1.37 ± 0.29 | <0.001 |

**Supplementary Table 2 - Mean vertex-wise explained variance of the four predictive models in the test dataset.** The vertex-wise explained variance values were averaged over vertices within the range of 1-8º of eccentricity within the dorsal portion of V1-3, early visual cortex, and higher order visual areas.

| | **POLAR ANGLE** | |
|---|---|---|
| | | **Mean $r^2$ ± std** |
| *Hemisphere* | *Areas* | *Model* |
| LEFT | Dorsal V1-3 | 0.21 ± 0.22 |
| | Early visual cortex | 0.19 ± 0.20 |
| | Higher order areas | 0.13 ± 0.15 |
| RIGHT | Dorsal V1-3 | 0.12 ± 0.13 |
| | Early visual cortex | 0.13 ± 0.15 |
| | Higher order areas | 0.14 ± 0.16 |
| | **ECCENTRICITY** | |
| | | **Mean $r^2$ ± std** |
| *Hemisphere* | *Areas* | *Model* |
| LEFT | Dorsal V1-3 | 0.17 ± 0.21 |
| | Early visual cortex | 0.15 ± 0.18 |
| | Higher order areas | 0.12 ± 0.15 |
| RIGHT | Dorsal V1-3 | 0.13 ± 0.15 |
| | Early visual cortex | 0.15 ± 0.15 |
| | Higher order areas | 0.13 ± 0.15 |

**Supplementary Table 3 – Results from the 3 X 3 repeated measures ANOVA performed to test for effects of Prediction Type (our model, the group-average map, and the Benson et al. (2014) model) and Empirical Dataset used (pRF fit 1, fit 2, and fit 3) on the prediction errors.** A main effect of Prediction Type was found ($p<0.05$).

| WITHIN SUBJECTS EFFECTS | | | | | |
|---|---|---|---|---|---|
| | **Sum of Squares** | **df** | **Mean square** | **F** | **p-value** |
| *Prediction Type* | 116.13 | 2 | 58.063 | 5.222 | 0.016 |
| *Residual* | 200.16 | 18 | 11.120 | | |
| *Empirical Dataset* | 4.26 | 2 | 2.129 | 0.855 | 0.442 |
| *Residual* | 44.82 | 18 | 2.490 | | |
| *Prediction Type × Empirical Dataset* | 2.65 | 4 | 0.661 | 1.938 | 0.125 |
| *Residual* | 12.29 | 36 | 0.341 | | |

**Supplementary Table 4 – Post hoc comparison with Bonferroni correction of the main effect of Prediction Type found in the 3x3 repeated measures ANOVA (Supplementary Table 3).** Our model's predictions were found to be significantly more accurate than group-average-based predictions ($p < 0.05$).

| COMPARISON | | | | | | |
|---|---|---|---|---|---|---|
| **Prediction** | **Prediction** | **Mean Difference (º)** | **SE** | **df** | **t** | $P_{bonferroni}$ |
| Our Model | - Benson et al. (2014) | -1.45 | 0.864 | 9.00 | -1.68 | 0.380 |
| | - Average | -2.78 | 0.612 | 9.00 | -4.54 | 0.004 |
| Benson et al. (2014) | - Average | -1.33 | 1.050 | 9.00 | -1.26 | 0.713 |

**Supplementary Table 5 – Retinotopic map models in perspective.**

| FEATURES | MODELS | | |
|---|---|---|---|
| | **Bayesian model** (***prior only***[1]) - Benson and Winawer, 2018 | **Bayesian model** (***prior + posterior***) - Benson and Winawer, 2018 | **DeepRetinotopy** - Ribeiro, Bollmann and Puckett, 2020 |
| **Input - Anatomical data** | ✓ | ✓ | ✓ |
| **Input - Functional data** | | ✓ | |
| **Dataset** | 8 participants | 8 participants | **181 participants** |
| **Polar angle error (V1-3)** | 23º | 25º[2] | **20.24º** |
| **Eccentricity error (V1-3)** | 1.08º | 0.76º[2] | **0.88º** |

[1] Note that the prior template map from the Bayesian model of retinotopic maps (Benson and Winawer, 2018) is an updated version of the template map in Benson et al. (2014), which in turn is an updated version of the retinotopic map template in Benson et al., (2012).
[2] Note that some differences exist in how the errors were calculated across the studies.

**Supplementary Table 6 - Participants' HCP identification number.** Identification number of the participants within the test and development datasets.

| DATASET | HCP ID | PARTICIPANT NUMBER |
|---|---|---|
| **Test** | 680957 | 1 |
| | 191841 | 2 |
| | 617748 | 3 |
| | 725751 | 4 |
| | 198653 | 5 |
| | 191336 | 6 |
| | 572045 | 7 |
| | 601127 | 8 |
| | 644246 | 9 |
| | 157336 | 10 |
| **Development** | 186949 | - |
| | 169747 | - |
| | 826353 | - |
| | 825048 | - |
| | 671855 | - |
| | 751550 | - |
| | 318637 | - |
| | 131722 | - |
| | 137128 | - |
| | 706040 | - |

**Supplementary methods**

In traditional deep learning, data is represented in a regular grid – i.e., adjacent pixels (or voxels) are regularly spaced from each other. Hence information about the organization of that grid-like structure is already embedded in deep learning architectures. For example, for object recognition problems, data are usually 2D images with a similar grid-like structure regardless of the dataset. Cortical surface models, however, are not regularly represented (having varying distances between vertices). To perform geometric deep learning with surface data then, the model must be provided with the representation the surface topology. Note that the position of the vertices in the 3D space and the connections among the vertices is used by the convolutional layer to enable information to be found and aggregated in local neighborhoods. In our model, curvature and myelin values (explicit features) as well as the 3D coordinates and connectivity of vertices (implicit features) are all used as input for the model. To better appreciate the data structure, an executable code notebook hosted on Google Collab is currently available that allows anyone to load the trained models, to generate predictions on the test dataset, and to check out the structure of data objects (https://colab.research.google.com/drive/1zXrK5HK806iXkJ6IIKeg0ePKm1SCTQlB?usp=sharing). A brief explanation of data structure is provided below.

Data objects consist of multiple attributes, such as X, Y, edge_index and edge_attr (for more information, see the PyTorch Geometric documentation at https://pytorch-geometric.readthedocs.io/en/latest/modules/data.html). 'Data.x' refers to vertices' features, myelin and curvature in our case, with dimension equal to [number of vertices, 2]. 'Data.y' refers to vertices' features that we aim to predict, polar angle or eccentricity, with dimension equal to [number of vertices, 1]. 'Data.edge_index' is a matrix with the indices of vertices that form edges, with dimension equal to [2, number of edges]. Finally, 'data.edge_attr' is a matrix with the relative distance between pairs of vertices forming edges, with dimension equal to [number of edges, 3]. Spline-based convolutional layers (SConv(k,Min,Mout); Fey et al., 2018) take as input 'data.x', 'data.edge_index' and 'data.edge_attr' to aggregate information in local neighborhoods by looking at a target vertex at a time. The topology of the convolutional layer's output is the same as the input, with only 'data.x' varying from one layer to the other. Therefore, in our model, the inner representations do not change in terms of number of vertices and their connections from one convolutional layer to the next. Number of vertices were equal to 3,267 for the left hemisphere and 3,219 vertices for right hemisphere, adding up to 19,024 edges for the left hemisphere and 18,760 edges for the right hemisphere.
